# Mirror Neurons are Modulated by Grip Force and Reward Expectation in the Sensorimotor Cortices (S1, M1, PMd, PMv)

**DOI:** 10.1101/2020.12.30.424850

**Authors:** Md Moin Uddin Atique, Joseph Thachil Francis

## Abstract

Mirror Neurons (MNs) respond similarly when primates make, or observe, grasping movements. Recent work indicates that reward expectation influences rostral M1 (rM1) during manual, observational, and Brain Machine Interface (BMI) reaching movements. Previous work showed MNs are modulated by subjective value. Here we expand on the above work utilizing two non-human primates (NHPs), one male *Macaca* Radiata (NHP S) and one female *Macaca* Mulatta (NHP P), that were trained to perform a cued reward level isometric grip-force task, where the NHPs had to apply visually cued grip-force to move and transport a virtual object. We found a population of (S1 area 1-2, rM1, PMd, PMv) units that significantly represented grip-force during manual and observational trials. We found the neural representation of visually cued force was similar during observational trials and manual trials for the same units, however, the representation was weaker during observational trials. Comparing changes in neural time lags between manual and observational tasks indicated that a subpopulation fit the standard MN definition of observational neural activity lagging the visual information. Neural activity in (S1 areas 1-2, rM1, PMd, PMv) significantly represented force and reward expectation. In summary, we present results indicating that sensorimotor cortices have MNs for visually cued force and value.

## Introduction

Mirror neuron (MN) activity has been observed while studying kinematic behaviors, ^1–4^, and through indirect measures in human motor cortex ^5^. In order to ask questions about force information at the single unit level, we utilized an isometric grip-force paradigm (Fig.1), and cued reward level ^6, 7^, while modulating both cued grip-force and reward level in the current work. We utilized this grip-force paradigm initially for brain machine interfacing research ^8^, and subsequently included observation only trials to ask questions about mirror neuron activity within the sensorimotor regions when kinematics are not actively being controlled, but rather force output is the controlled variable.

**Figure 1:**
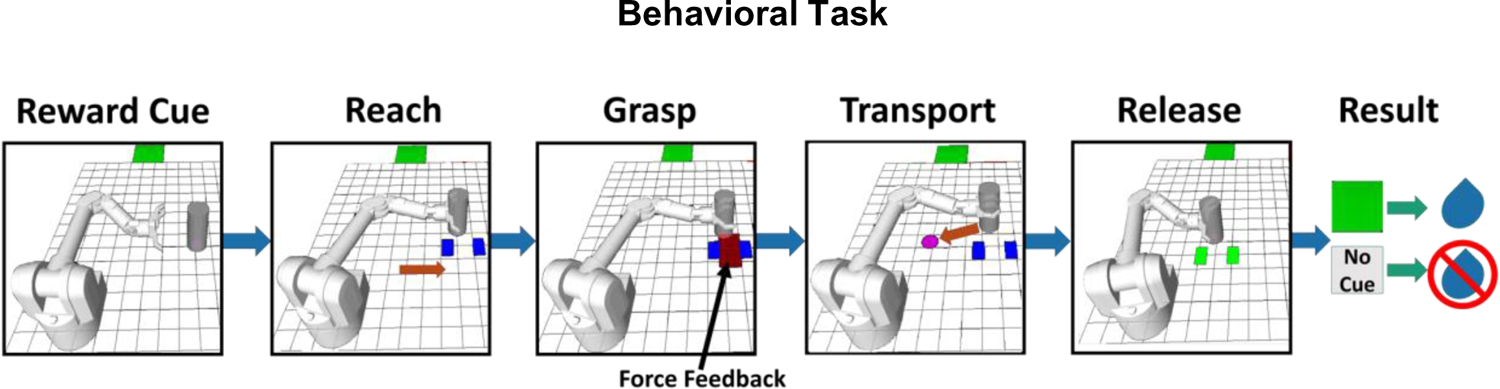
Flow diagram illustrating a single trial. The reward cue (green square, R1) was shown for R1 trials in cued blocks only. Absence of the green square indicated a non-rewarding (R0) trial during cued blocks of the task. The robotic arm started in a resting position for all trials (left-most image). During uncued blocks, there was no cue provided during the “Reward cue” period for any trials. Please note, under experimental conditions with NHPs the background of the grid and free space around the robotic arm and object were black. For the reader’s benefit and visual clarity, both areas have been changed to white here. Blue rectangles are grip-force targets and were seen for the duration of a trial. Red rectangle shows visual force feedback of the NHPs grip-force output initially but is no longer shown once the NHP has reached the acceptable force range (blue rectangles).

A better understanding of how reward expectation influences the primary- and pre-sensorimotor cortices has practical implications towards the production of stable and autonomously updating brain machine interfaces (BMIs) ^6, 7, 9–12^. Neural modulation related to reward has been well characterized within many brain structures ^13–15^, and is known to occur at many levels, such as single units and local field potentials (LFP) within rM1 ^6^. The neural response to both reward and conditioned stimuli (CS) that predict reward have been demonstrated in a multitude of brain regions ^7, 16–22^. It has recently been shown that reward expectation changes directional and force related tuning functions within the primary sensorimotor cortices during reaching movements and BMI reaching movements ^8, 23^. The above studies indicate the need to conduct more research on the influence reward has on these brain regions, and how such variables may modulate MNs. The influence of reward on the caudal primary somatosensory cortex (cS1, area1 and 2) was reported in conference proceedings ^24–26^. While these experiments demonstrated our ability to classify reward expectation from S1 they did not provide insight on its influence of grip-force during action and action observation as we do here. A similar reward signal has been demonstrated in primary somatosensory cortex utilizing fMRI, where a stronger BOLD response was recorded for higher reward delivery ^27^, the investigation of reward-correlated signals within S1 at the single unit level has not been fully developed.

Our previous work focused on identifying significant units related to reward and/or force within single blocks of data ^8^ during manual trials and BMI control. To expand on these findings, we thought it important to investigate the cause behind each unit’s modulated activity across multiple blocks, during both manual ^8^, and observational tasks. We hypothesized modulation responses during observation might be explained by the presence of mirror neural activity when NHPs observed the task, received reward, or possibly both. To this end, our goal was to detect units that encoded the following: 1) Motor actions that are the physical application of force; 2) Cued “force” during visual observation; 3) units that encoded both 1, and 2. 4) Determine which of these (1-3) were further modulated by reward level, cued or uncued. We focused on “grasping” movements with cued isometric grip-force control, to expand beyond our previous work on reaching movements ^6, 28^. Below we demonstrate our ability to decode actual and observed grip-force neural responses, during manual and observational trials, as well as reward’s influence on the neural population during reward cued and uncued trials. Others have described activity of MNs in rM1 during observation of movement ^3, 28–31^, but little has been reported on the activity in S1 during observational tasks, or any of the 4 regions as it pertains to isometric grip-force control while modulating reward level as we do here. There are some studies on human and NHP that showed evidence of MN activation ^32–34^ and grip force related modulation ^35, 36^ is present in somatosensory areas which implies the possibility of MN related activity due to grip force in cS1 cortex. Generally, our paper presents findings of MN responses related to varying levels of grip-force (Fig.6) and reward’s influence on grip-force MN activity (Fig.8-10) within cS1, rM1, PMd and PMv.

## Methods

All NHP manipulations described in this work were approved by the Institutional Animal Care and Use Committee of the State University of New York at Downstate Medical Center and conformed to National Institutes of Health (NIH) and United States Department of Agriculture (USDA) animal care and use guidelines. In addition, this work complies with the ARRIVE guidelines. Two non-human primates (NHPs), one 9.0kg male *Macaca* Radiata (NHP S) and one 5.0kg female *Macaca* Mulatta (NHP P), were trained to perform a behavioral grip-force task using their right hand and subsequently implanted with 96-channel electrode arrays in cS1, rM1, PMd and PMv.

We describe the sequence for one complete trial of the manual reward-cued version of the task seen in Fig.1. First, the virtual robot moves to the start position, which is always to the far left of the task space. Second, cue scene, a green square moved across the top of the screen for 0.5s from the left to indicate a rewarding (R1) trial, absence of this green square indicated a non-rewarding (R0) trial. Third, the virtual robot autonomously moved its arm toward the cylindrical object in the reach scene. Fourth, during the grasp scene of manual trials, the NHPs had to apply isometric grip-force using a stationary handle that contained a force transducer. The amount of force required was indicated by two blue rectangles, where the inner vertical edge of each blue rectangle indicated minimum force, and the outer edges indicated maximum force. The minimum threshold of force for NHP S was randomly chosen from either 150 or 200 and for NHP P it was 100 or 150. If the force output went below that minimum value the trial was considered a failure. The upper threshold was determined by randomly adding either 300 or 500 to the minimum for NHP S and adding randomly either 250 or 350 to the minimum for NHP P. Therefore, the ranges used for NHP S were randomly selected from the following, 150-450, 150-650, 200-500, 200-700, and for NHP P the ranges were, 100-350, 100-450, 150-400, 150-500. If the output force of the NHP was above these max levels the trial was considered a failure. The peak grip force values across the manual trials, R1 and R0, in cued and uncued blocks are provided in supplementary figure S7. When the NHP’s applied force was within the acceptable tolerance range, the red force rectangle’s edges would also be within the blue rectangles. When this occurred, the visual force feedback (red rectangle) was removed, and the robot hand would grasp the object. We utilized this method so that the NHPs could not simply use visual feedback to perform the task, but rather had to learn to control their force output based on the visual force targets and somatosensory feedback for manual trials. Thus, they had to learn to produce a given grip-force, and this may have led to the positive results during observational tasks, as they most likely had built an internal representation of the force cues and their production of such force as well as the expected somatosensory feedback. During the transport scene, the robot arm autonomously moved the object to a predetermined target location, indicated by a pink circle. If the NHPs applied the appropriate grip-force during object grasp and transport, and then released when the object was touching the virtual ground at the pink target location, the trial was successful. Feedback was provided to NHPs for successful placement when the blue force squares turned green, and the robotic arm released the cylinder during the release scene. If they did not apply proper force during initial grasp, failed to maintain an acceptable force range at any time during the grasp or transport scenes, or released their grip early or late, the trial was considered a failure.

There were four types of task blocks, which were experienced by each NHP in the following order: first, manually performed tasks with the presence of a reward level conditioned stimulus (CS) (manual cued); second, manually performed tasks without a reward level CS (manual uncued); third, observational task with the presence of a reward level CS (observational cued); fourth, observational task without a CS (observational uncued). All trials (R0 and R1) during uncued blocks lacked a visual reward cue so the NHP had no indication of trial value until the post result period, and thus, no explicitly cued expectation for reward outcome as the trial sequence was randomized with no clear autocorrelation (see supplementary Fig.S1). The time limit to complete a successful trial was within 10s. The NHPs had to repeat failed trials under the same reward conditions until successful, thus giving incentive to perform non-rewarding trials. Without this added stipulation the NHPs would choose to fail R0 trials to move onto a possible R1 trial, indicating the NHPs clearly understood the cue reward values. Juice rewards were delivered via a system-controlled solenoid driven by task logic (Crist instruments). All elements of the grip-force task were developed in Linux using robot operating system (ROS) ^37^. ROS and Python controlled the task logic, outputs to the reward delivery system, and provided timestamp synchronization with external systems to simultaneously and accurately record task state, reward delivery, and neural data.

During observational blocks NHPs were visually monitored by researchers in real-time via cameras during all sessions to make sure NHPs remained attentive and focused on the projection screen, especially during observational trials. During observational trials NHPs did not have access to the force transducer handle, and their arms were blocked behind a plexiglass box meant to keep the NHPs hands away from the trainers (BKIN’s Arms-Free restraint chair). Additionally, NHPs were trained, and performed all experiments in a dark, quiet, and distraction-free isolation chamber to encourage their attention remained on the large projection screen. Each NHPs’ rear-mounted cranial head post was affixed to the BKIN primate chair to restrict head movement for neural recordings. The virtual environment was projected onto a vertical screen in the animal’s visual field. The visual projection of the virtual robotic system was approximately the same size as the real WAM robotic arm ∼1m reach (WAM Barrett). Recording sessions were broken into blocks of trials that averaged ten minutes of randomized R1 and R0 trials.

### Chronic Implantation

We performed implantation procedures as described in-depth in a previous methods paper ^38^, but give a summary here. Following training, when both NHPs achieved greater than 80% success rate on all trials in a block, the animals were implanted with chronic electrode arrays consisting of a 10 by 10 array of 1.5mm electrodes of which 96 were active for all regions (rM1, PMd, PMv) except cS1, which was implanted with electrodes of 1mm length. Spacing between individual electrodes within the arrays were 400μm (Utah array, Blackrock Microsystems) (Fig.2). All surgical procedures were conducted using aseptic technique. NHPs were initially anesthetized with Ketamine, followed by isoflurane and a continuous infusion of fentanyl. The NHP was then placed into a stereotactic frame before the surgical site was shaved and cleaned. An incision was made along the skull to expose desired implant locations. The craniotomy window was large enough to accommodate implant locations while leaving enough margin between the dural flap and skull. The dural flap was kept under tension using stay sutures until electrode arrays were implanted and the site was ready to close. We performed intraoperative probing of cS1 to ensure implantation within the hand region, by using a four shank, 32-channel silicon microelectrode array ^39^ within the post-central gyrus as determined by stereotactic coordinates. The NHP’s contralateral hand was continuously stimulated by touch to assess cS1 hand region boundaries. Neural responses were amplified and sent to an audio speaker to verify stimulation areas. Utah arrays were then chronically implanted in cS1’s hand region, in rM1 directly reflected across the central sulcus rostral from cS1 for NHP S. However, NHP P had large blood vessels that interfered, and we had to implant rM1 more lateral than in NHP S as seen in Fig.2. In addition, as seen in Fig.2, NHP P’s PMd implant was slightly more medial than NHP P, and NHP S’s PMv was slightly more lateral than NHP P. In both NHPs our PMv arrays had fewer clear single units than the other arrays. We cannot be certain but believe this was due to the PMv wire bundles exiting the craniotomy closer to the array and at a stiffer region of the wire than the other arrays. The craniotomy was replaced according to previously described methods ^38^.

**Figure 2:**
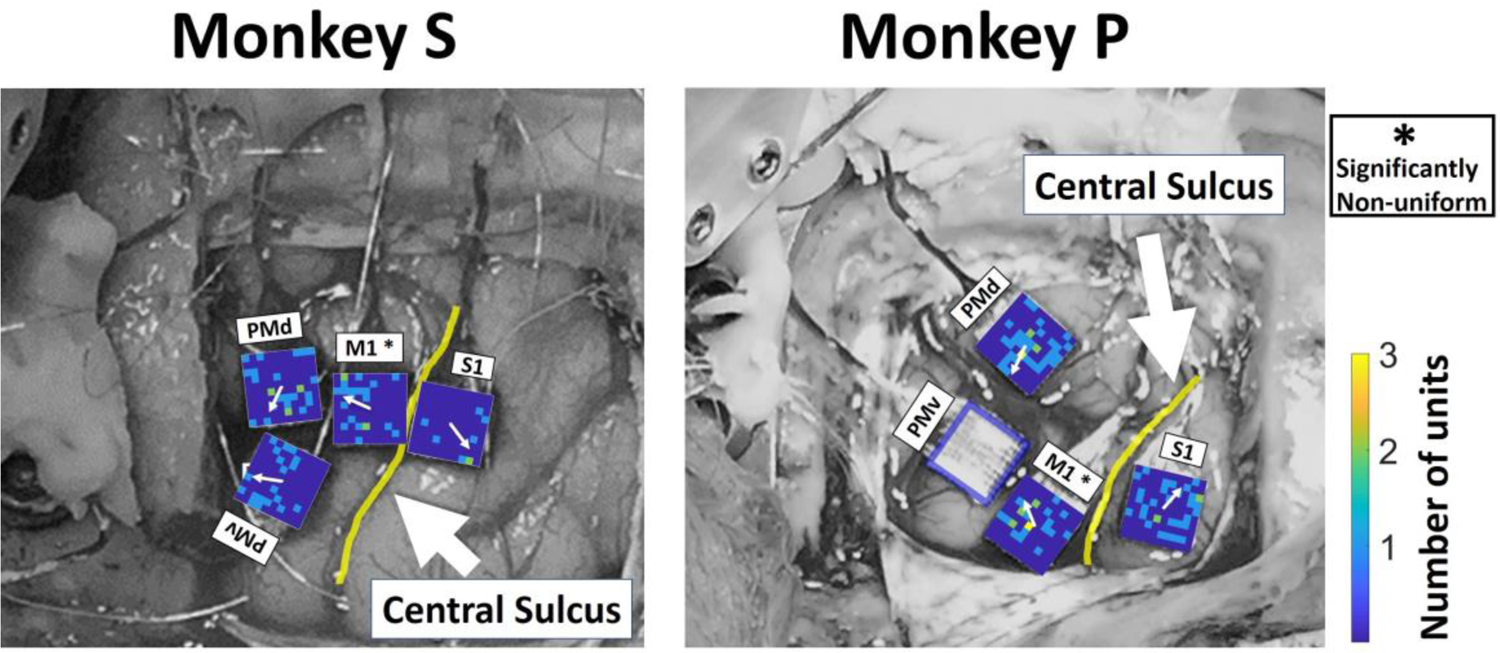
Position of four Utah arrays in relation to the central sulcus for NHP S (left) and P (right). The four arrays were implanted in cS1, rM1, PMd and PMv cortices. The yellow line indicates the Central Sulcus. Note in NHP P we had to implant the rM1 array more lateral than in NHP S due to a large set of blood vessel running through that region. Likewise, PMd was implanted more medial in NHP P for this same reason as compared to NHP S. The color map on each array is for the number of MN (mirror neuron) units recorded in each location, we did not have the electrode map for NHP P PMv. The white arrow inside the color map represents the mean direction of MN position with respect to the center of the array. A Rayleigh test (p<0.01) showed the distribution of MNs around the center of the array is non-uniform only for M1 in both NHP.

### Neural Recording

Neural recordings were performed using three synchronized multi-channel acquisition processors (Plexon, Dallas, TX), each having 128-spike waveform recording channels and 32 analogue channels used for simultaneous recording of local field potentials ^6, 12^. Single unit recordings were amplified and retained using waveform voltage thresholds. Thresholds were set using an auto-scale (Plexon recording software) feature, followed by manual adjustments that eliminated noise on channels prior to recording. Robot operating system ^37^ and Python programs controlled task logic and embedded timestamps into neural recordings using a common clock. The common clock was maintained by a microprocessor that delivered a 2kHz pulse to keep task logic and neural data synchronized. Initially, unit waveforms were automatically clustered using a k-means algorithm in principal component space ^40, 41^. Afterward, we used Plexon’s offline sorter software to adjust clusters to remove noise and artifacts.

### Significance Related to Reward

We only considered successfully completed trials for all analyses and results presented in this work. We analyzed single unit activity from four types of task blocks for each NHP, cued or uncued, and these were either manual or observational. For trials that contained a visual reward cue, the post-cue (0-500ms) analysis window began immediately after the green visual cue came to rest. In uncued trials, the “post-cue” period (0-500ms) began when the robotic arm returned to its rest position from the previous trial. We defined the “post-result” period (0-500ms) as the time after the cylindrical object was successfully placed at the target location. We analyzed spike activity in the post-cue and post-result periods separately for cued and uncued blocks to identify units significant for reward modulation. We binned the spike activity into non-overlapping bins of 100ms covering 500ms for both the post-cue and post-result periods. We collected the p-values from t-test for Spearman Rank correlation and the test was done between the binned spike rate and corresponding reward levels (0 for R0 and 1 for R1). For reward levels, 0 is used for R0 and 1 is for R1. We then extracted units that were significant (t-test, p <0.05) for reward expectation (post-cue activity, R1 vs. R0 trials) and reward result (post-result activity, R1 (after reward delivery) vs. R0 (after successful trial with no reward)). The p-values from the test are adjusted for multiple comparisons using the false discovery rate (FDR) procedure by Benjamini and Hochberg (BH Method) (Benjamini et al. 1995) for number of times t-test was applied which is equal to the number of units for each case, which is given in supplementary (Fig.S2). Finally, we compared spike rate activity between cued and uncued blocks to gain better insight as to how reward feedback information was affected by the presence or absence of a reward cue that is reward expectation based on explicit environmental cues. Our interest in reward expectation stems from our work towards autonomously updating BMIs, and thus our use of this paradigm.

### Grip-force Trajectory Prediction

We applied linear regression in two steps to identify units that showed significant prediction of grip-force from cS1, rM1, PMd and PMv. In these two steps, the first step identified all units that were significantly related to grip-force, which was used below in the sections labeled “*Grip-force Tuning Curve Analysis”* and “*Identifying Observation Modulated Neurons*”. The second step was applied to sort out a selective number of units, from the units collected in the first step, for force prediction to reduce the number of predictors and avoid computational complexity. Unit activity from the aforementioned cortices were analyzed along with force profiles for both NHPs during manual and observation task blocks, see supplementary material for data not shown in the main text. During some trials, initially the NHP’s applied more grip-force than necessary and in an effort to meet the target value, reduced their grip-force drastically. This resulted in an overcorrection where the NHP had to apply more force to meet the requirement and resulted in force profiles with multiple peaks. We manually inspected and pruned all force profiles that contained multiple peaks due to highly variable or corrective grip-forces applied by NHPs. We defined force onset as the point where force values increased from zero to positive value of 50 au and force offset when values returned below 50 au, as the NHPs only grasped the force transducer during the grasp scene, and a trial was considered successful when their output force < 50 au, which generally meant they let go of the handle. Force data was collected from 0.5s prior to force onset through 0.5s after force offset for all trials. The collected force data was smoothed using a Gaussian kernel (100ms wide). During observational trials the visually cued “force” targets, and “force” output profiles were trapezoidal and were inferred from the trapezoidal velocity grasping motion of the hand. Therefore, smoothening gave the “force” profile a gaussian shape like the force data on manual blocks. From the force data collected in each trial, we evaluated neural spike activity 0.5s prior to, and following each force sample value. This neural activity was further placed into ten non-overlapping time bins centered on each force value (100ms bins, covering 0.5s pre- and 0.5s post-force sample value) to determine a unit’s significance to grip-force, so for each unit there were 10-time bins used as predictors. We performed this separately on each unit from all four cortices. As we smoothed the unit raw data using a Gaussian kernel (100ms wide), we wanted to find a generalized kernel width that would be used for both cued and uncued data blocks of the same kind, manual vs. observational. The force prediction accuracy for different kernel bin widths from 10ms to 250ms were determined for all data. We manually selected the above value (100ms) for the kernel width so that the predictions were close to the peak prediction values for all blocks. See Fig.S3 for differences between analysis on the smoothed data, vs. unsmoothed. The collected binned data was square root transformed to achieve a more normalized distribution. To make sure this square root transformation is not affecting the prediction results significantly a comparison of grip force prediction accuracy is given before and after the transformation is applied in supplementary Fig.S4.

Data recorded from each NHP was split into a fit (90%) and test (10%) set to check goodness of fit and verify predictive performance of the final linear regression model (equation 2). However, before testing the final linear regression model, we extracted significant units related to force in a two-step process. To do this, we subdivided the 90% fit set into 80% (fit) and 10% (validate) data subsets, see the supplementary section for a cartoon of this procedure Fig.S5.

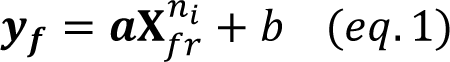

Equation 1 shows the linear model used, where *y*_f_ is the vector of force values, *X^ni^_fr_* is the binned spike rate matrix for neuron n_i_, while *a* and *b* are coefficients fit to the data. In the first step, we applied linear regression using equation 1 with the 80% subset data for the fit and collected the F-statistics from the analysis of variance (ANOVA) for the model. Units were sorted according to their p-value from the above F-test and only those with a significant fit (p <0.05) were considered in the second step of the process. For the second step, we took significant force units, starting with the most significant (lowest p-value), and used it to test prediction of force values in the remaining 10% validation subset. We calculated the R-squared value after each unit was added to the model and if the R-squared value increased (improved prediction results), the unit was kept. Remaining units that did not improve the model’s force prediction were pruned. After the two-step process, the subset of significant force units was utilized in the final linear regression model (eq. 2). This subset of units was used with updated coefficients fit to the original 90% validation dataset. Finally, we used the held out 10% test set to determine prediction and validate accuracy of the linear regression model (eq. 2) by comparing them against actual recorded force values.

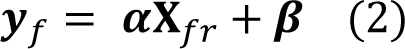

The regression model used for final grip-force prediction from neural activity is shown in equation 2. Here *y*_f_ denotes grip-force and *X*_fr_ represents binned firing rates for the population of units being used, *α* and *β* are model coefficients fit to the data.

### Grip-force Tuning Curve Analysis

Our previous research concerning force tuning curves in rM1 showed a significant difference between R1 and R0 trials ^8^. We applied the same analysis to our cS1, rM1, PMd and PMv data to determine if neural activity led to different force tuning curves when taken from R1 vs. R0 trials. As in Zhao et al. 2018 we utilized analysis of covariance (ANCOVA). We analyzed each unit previously identified as significant for force to measure whether the slopes of the tuning curves between R1 and R0 were significantly different (F-test, p <0.01). The p-values from the ANCOVA test are adjusted for multiple comparisons using the false discovery rate (FDR) procedure by Benjamini and Hochberg (BH Method) (Benjamini et al. 1995) for number of population on which the test was applied. Significant units identified by ANCOVA were considered to have a force representation that was modulated by reward expectation, and units that also passed the BH method are stated explicitly.

### Identifying Observation Modulated Neurons

We started by identifying and tracking single unit activity across multiple blocks of recorded data. We compared single unit activity between reward-cued manual and observational blocks, and again between reward-uncued manual and observational blocks for each NHP and cortices (CS1, rM1, PMd and PMv). We tracked single unit activity from the two manual and two observation blocks performed in a single day for each NHP. If a unit on a channel retained the same waveform over all 4 recorded blocks, we considered it the same unit. In addition, we verified this single unit activity by checking the correlation coefficient between the waveform shapes across all blocks, where a high correlation (> 0.98) indicated the same unit. We also confirmed the consistency of single unit activity across blocks by checking the first two principal components using principal component analysis (PCA). Performing these checks allowed us to track single unit activity with significant correlation to reward, force, or both during manual and observation task during cued and uncued blocks towards identifying putative mirror neural activity responding to force and reward modulation.

## Results

The results section is structured as follows: 1) We start by presenting raw data from manual and observational versions of our isometric grip-force task showing peri-event-time-histograms (PETHs) and rasters of sample units, indicating the modulation of single units by both cued grip-force and cued reward level during manual and observational trials in the caudal S1 areas 1-2 (cS1), rostral M1 (rM1), PMd and PMv, for NHP S and P, Fig.3 and Fig.4 respectively. 2) We present population results from linear regression focusing on cued grip-force, showing that subpopulations of these brain regions encode grip-force trajectories (Fig.5) even during observational trials. 3) In Figure 6 we test for temporal shifts between the neural representation of force, between manual and observational versions of the tasks, to determine if the same units are acting in response to the visual input, classical MNs, or predicting such input, as expected by mental simulation of the predicted visual stimulus, which could in turn be either mental simulation of the expected movement ^2^, or MNs responding in an anticipatory way due to the familiarity with the task ^43^. 4) Subsequently, we give information on units that are modulated by both cued grip-force and cued reward level during the force output period of the manual and observational tasks for the same single units that is putative MNs for force modulated by reward. To be clear, the term force, when used for the observational trials, indicates the force that would be expected from the NHP if the NHP were performing the task manually based on the visual force target cues still being presented during the observational task.

**Figure 3:**
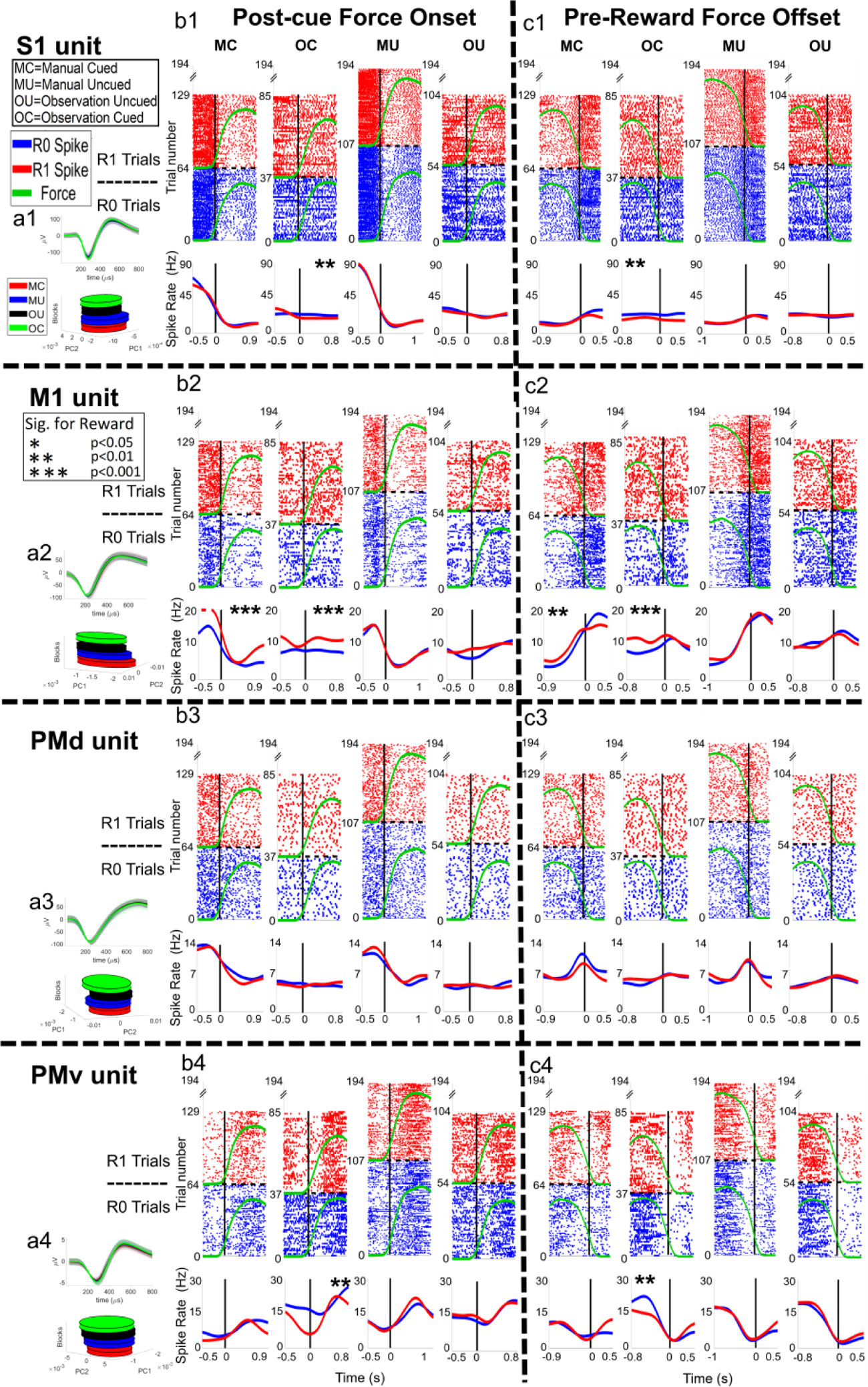
Raster plots of example units from NHP S during the manual and observational blocks. The plot numbers, 1-4, indicate the brain regions (1-4 for cS1, rM1, PMd and PMv respectively). The letters, a-c, after the numbers, 1-4, are associated with (a) single unit waveform for the example unit, (b) force onset plots and (c) for force offset plots. For (b) and (c) there are four plots for manual cued task (MC), observational cued task (OC), manual uncued task (MU), and observational uncued task (OU) sequentially. The post-force onset and pre-force offset time is set as 60% of the mean force length of all trials for that task. On each subplot (b, c), the x-axis represents time in seconds, for raster plots the y-axis represents trial number, for spike rate plots (bottom of subplots) y-axis is spike rate in ‘Hz’. The dashed horizontal black line on each plot divides the R1 trials from R0 trials. Below the raster plots, solid red and blue lines indicate mean spike rate (Hz) for R1 and R0, respectively. An asterisk (*) indicates post force onset or pre-force offset spike activity is significantly (t-test, p<0.05, p<0.01 and p<0.001 denoted by *, ** and *** respectively) different between R0 and R1 trials.

**Figure 4:**
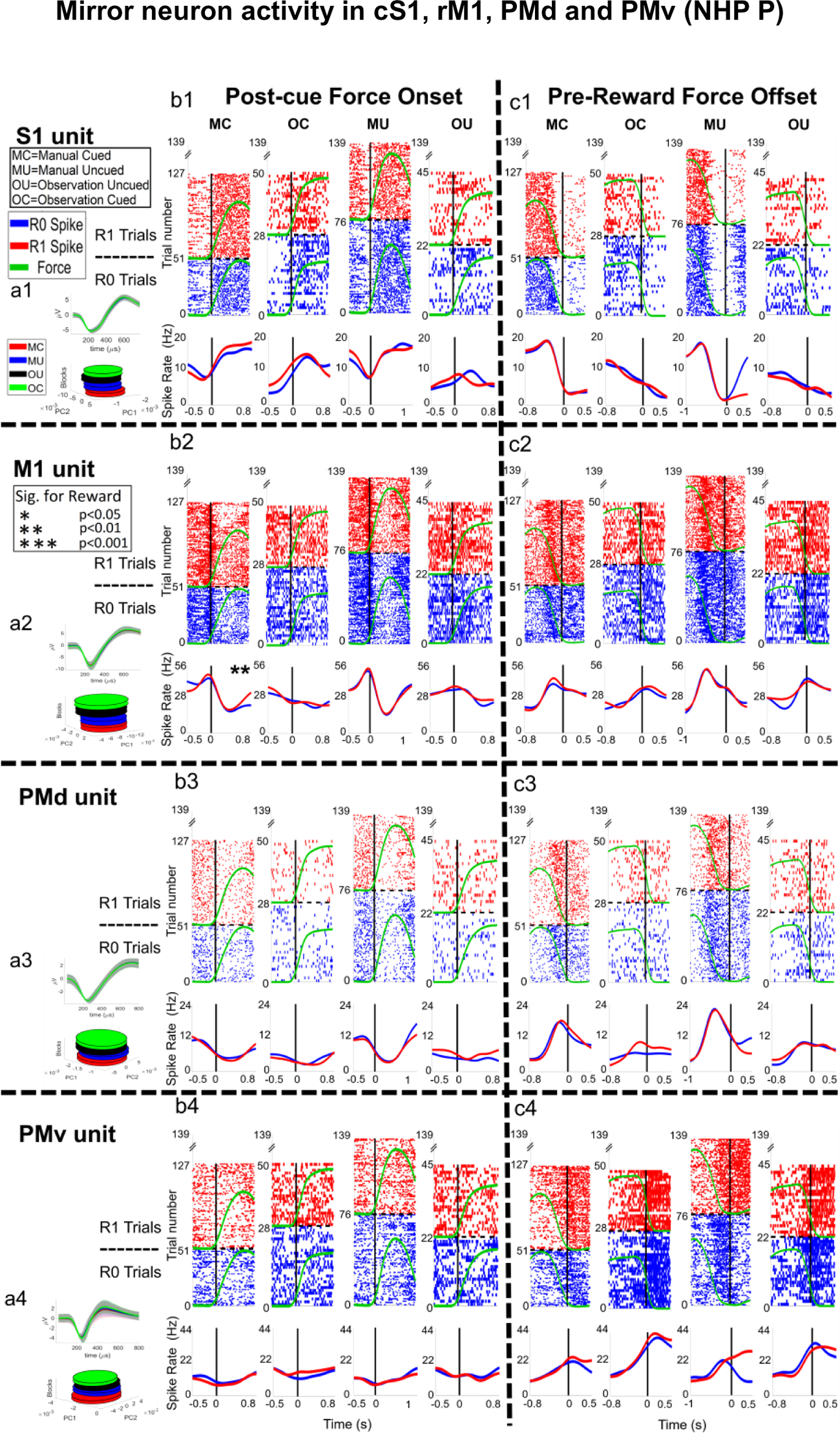
Raster plots of example units from NHP P during the manual and observational blocks. The plot numbers, 1-4, indicate the brain regions (1-4 for cS1, rM1, PMd and PMv respectively). The letters, a-c, after the numbers, 1-4, are associated with the (a) single unit waveform for an example unit, (b) force onset plots and (c) force offset plots. For (b) and (c) there are four plots for, manual cued task (MC), observational cued task (OC), manual uncued task (MU), and observational uncued task (OU) sequentially. The post-force onset and pre-force offset time is set as 60% of the mean force length of all trials for that task. On each subplot (b, c), the x-axis represents time in seconds, for raster plots the y-axis represents trial number, for spike rate plots (bottom of subplots) y-axis is spike rate in ‘Hz’. The dashed horizontal black line on each plot divides the R1 trials from R0 trials. Below the raster plots, solid red and blue lines indicate mean spike rate (Hz) for R1 and R0, respectively. An asterisk (*) indicates post force onset or pre-force offset spike activity is significantly (t-test, p<0.05, p<0.01 denoted by * and ** respectively) different between R0 and R1 trials.

**Figure 5:**
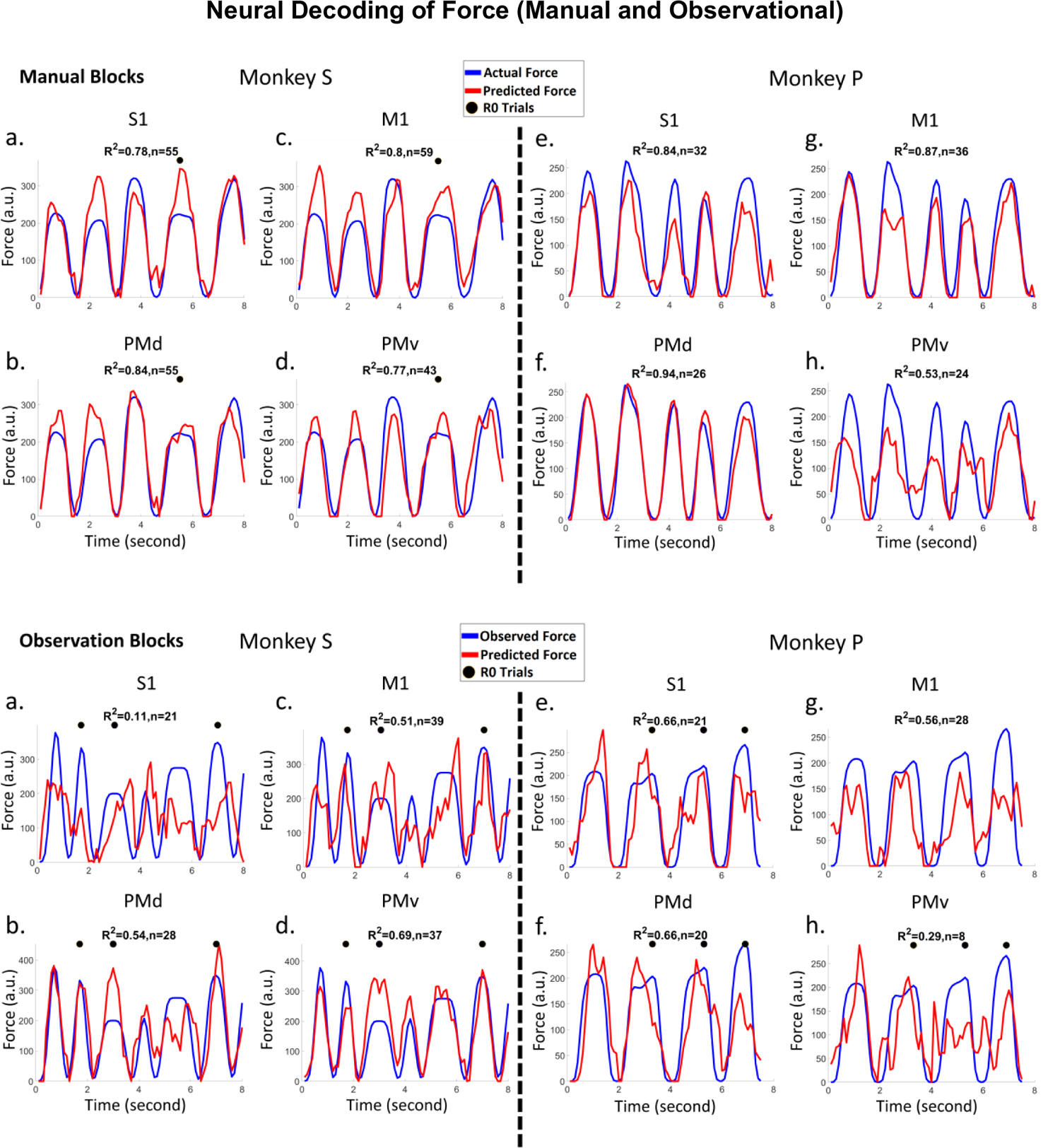
Force decoding (Red lines) from subpopulations of cS1, rM1, PMd, and PMv cortical recordings taken from manually performed blocks top two rows, and observational blocks bottom two rows. Linear regression (eq. 2) was used to predict the force from a subpopulation (see methods). The black dots on each plot indicate that force profile was taken from an R0 trial, while all others were R1 trials. Plots *a, b, c* and d show results for NHP S, for cS1, rM1, PMd and PMv cortices respectively. Plots e*, f, g,* and h show similar results for NHP P. The top of each figure shows R-squared values (R^2^) using the 10% test set and the number (n) of significant units used for regression to predict force. See table S2 for full stats on the regression models.

In Fig.3 and 4 (b-c) we present raster plots for example units with all trials aligned to the onset or offset of the grip-force. The mean force profile (indicated with green lines on the raster plots) shows force onset and offset, plotted with arbitrary units for R0 (blue raster) and R1 (red raster) trials. The mean duration that NHP S applied grip-force was 1.4 ± 0.4s (cued), 1.42 ±0.42s (uncued) during manual blocks. During observational blocks the “force” times where 1.12 ±0.26s (cued), 1.16±0.27s (uncued). For NHP P, the mean force duration was 1.2 ±0.28s (cued), 1.45 ±0.29s (uncued) during manual blocks, and 1.12 ±0.18s (cued), 1.21 ±0.23s (uncued) during observational blocks, see Fig.S6 – S8 for more on force level, duration and reaction times. NHP S achieved success rates of 77% (cued manual) and 82% (uncued manual) during the two manual-task blocks, while NHP P achieved rates of 58% (cued manual) and 72% (uncued manual). The number of R0 and R1 trials recorded on each individual block type is given in supplementary Fig.S9. The R0 and R1 mean force profiles are plotted with the same scale for comparison. All units shown in both Fig.3 (4 units) and 4 (4 units), where significant (F-test, BH corrected for p=0.05 for the number of units within the given brain region) for grip-force. For each NHP and cortices the number of units recorded is given in supplementary (Fig. S2), min ∼50 units and max ∼170 units. In Fig.3 ax (x=1, 2, 3 and, 4) show the mean spike waveform with standard error (shaded) for a single unit (top) and its PCA space (bottom) for all 4 blocks, manual cued (MC), manual uncued (MU), observational uncued (OU) and observational cued (OC), which were all recorded on the same day and in the aforementioned order for each NHP. Each raster subplot (b-e) shows activity for force onset (left) and force offset (right) periods.

Units in Figs.3 - 4 were chosen as they represented the variety of responses we saw in the population. We utilized 100ms bins during the post-force onset spike activity from the point when force sensor values crossed above 50 (a.u.) and pre-force offset activity until the reading went below 50 (a.u.). Fifty was the same value used for the task logic and was chosen based on experience with these NHPs performing this task in order to minimize false starts without missing movements. We collected p-values from t-tests on Spearman Rank correlation between the binned (100ms) spike rate and corresponding reward levels (1 for R0 and 2 for R1) indicating significance with an asterisk (*). For example, in Fig.3.b1-c1 we see an cS1 unit that decreases its activity at or just after the onset of “force” and is suppressed around the force offset period. Our hypothesis is this unit along with other example units that are showing suppression of activity is due to these units possibly being connected to extensor muscle groups of forearm and thus showing inhibitory spike activity during gripping action, see Fig.S17 for support of this. In addition, the unit in Fig.3.b1-c1 shows modulation by reward. Similarly, in Fig.3.b2-c2, we see an rM1 unit with a similar response to the cS1 unit, where the rM1 unit is suppressed at force onset and activated at force offset with some reward modulation as well. In Fig.3.b3-c3 a PMd unit is shown that is suppressed before force onset and has its peak activation pre-offset. Monkey P data was not as responsive to reward activity utilizing the spearman rank test as Monkey S. The Fig.3 responses shown for cS1, rM1 and PMd resemble extensor patters of activation while the PMv response shown resembles a more flexor typical response ^44, 45^. In Fig.4 we have chosen units that show some other responses seen in the population for both NHPs. The observational response to “force” was weaker compared to the actual force during manual trials and did not always align with the force onset and offset times as precisely. For additional raster plots on example units during grip force observation please see supplementary Fig.S10. All units seen in Figs.3-4 had significant fits to grip-force (F-test, p <0.05). This F-test was performed on the regression model (equation 1) to test whether it was a better fit than a degenerate model, which consisted of only a constant term. Figs. 3-4 show a total of 8 example units; 1 from each cortical region (rM1, cS1, PMd and PMv) for each NHP, P and S, from reward cued, and uncued blocks.

### Grip-force Decoding During Manual and Observational Trials

As described in the methods section and further in the supplementary section (Fig.S5), data blocks were analyzed using a two-step process. We identified significant units related to force using the simple linear model eq.1, *y*_f_ = *aX*^ni^_fr_ + *b*, where *y*_f_ is the vector of grip-force trajectory, *a* is the vector of regression parameters that multiply unit n_i_’s firing rate, where we used the associated F-statistic with a (p < 0.05) for significance determination. Note, we utilized neural data around a given grip-force timepoint, from −0.5s to 0.5s in 100ms bins for this regression, thus 10 bins. We extracted single units with activity that improved force decoding. Model 1 variables *a* (10 per unit) and *b* were fit using least-squares estimates from 90% of the recorded data (see Fig.S5 for data split cartoon). The remaining 10% of the data was used as a test set to verify the performance of force decoding. The R-squared values for decoding on the test set of force are shown below. We found high prediction accuracies for both NHPs. The results shown in Fig.5 are force decoding prediction from cS1, rM1, PMd and PMv cortices for cued manual blocks, top two rows, with the corresponding observational blocks seen in the bottom two rows. Note intertrial intervals have been clipped out for presentation purposes.

It is clear the manual tasks generally had higher levels of prediction as compared to the observational versions. We detected the units that were consistently present in all four data blocks (two manual and two observational) and those significant (F-test, BH adjusted for p=0.05 and number of units in the population) for grip-force fit in all data blocks. For NHP S we found 31 units in cS1, 42 in rM1, 46 in PMd and 29 in PMv that were consistently present in all four data blocks. For NHP P the numbers were 107 in cS1, 95 in rM1, 79 in PMd and 49 in PMv. Below in Fig. 6 we show the total percentage of these units from each region that showed significant linear regression model fit under both the manual and observational versions of the task that are the putative MNs. We used the peak value from the absolute correlation coefficients between grip force and each of the 10 bins of spike rate (100ms from −0.5ms pre grip force spike rate to 0.5s post, described “Grip force trajectory prediction” section inside method) to detect inhibitory or excitatory activity. We considered a unit’s representation excitatory if the actual value of that peak absolute correlation was positive otherwise it was considered inhibitory. We detected congruent units among the significant units that showed similar activity in all four data blocks of either inhibitory (blue) or excitatory (red) spike rate in relation to grip-force. Our hypothesis is the units that are showing inhibitory activity during grip force application are connected to the extensor muscle groups of the forearm which relaxes during gripping activity and show evidence for this in Fig.S17. Incongruent units showed opposing behavior between manual and observational tasks, such as excitatory during manual and inhibitory during observational, purple, or inhibitory during manual and excitatory during observational (green) in Fig.6. Other (gray) units didn’t follow any simple pattern during all four data blocks as seen in Figs.S11-S12. The number of units for each possible combination of inhibitory or excitatory spike activity during grip force for four data blocks are given in supplementary (Fig.S11 for NHP S and Fig.S12 for NHP P) section. Monkey S showed what one might expect PMv>PMd>rM1>cS1, whereas NHP P’s data was not in this clear expected order and could be due to the placement of the electrode arrays due to the vasculature-imposed restrictions as seen in Fig.2. However, in both NHPs the largest single group of MNs appear to be congruent and inhibitory for our task (blue bar sections).

**Figure 6:**
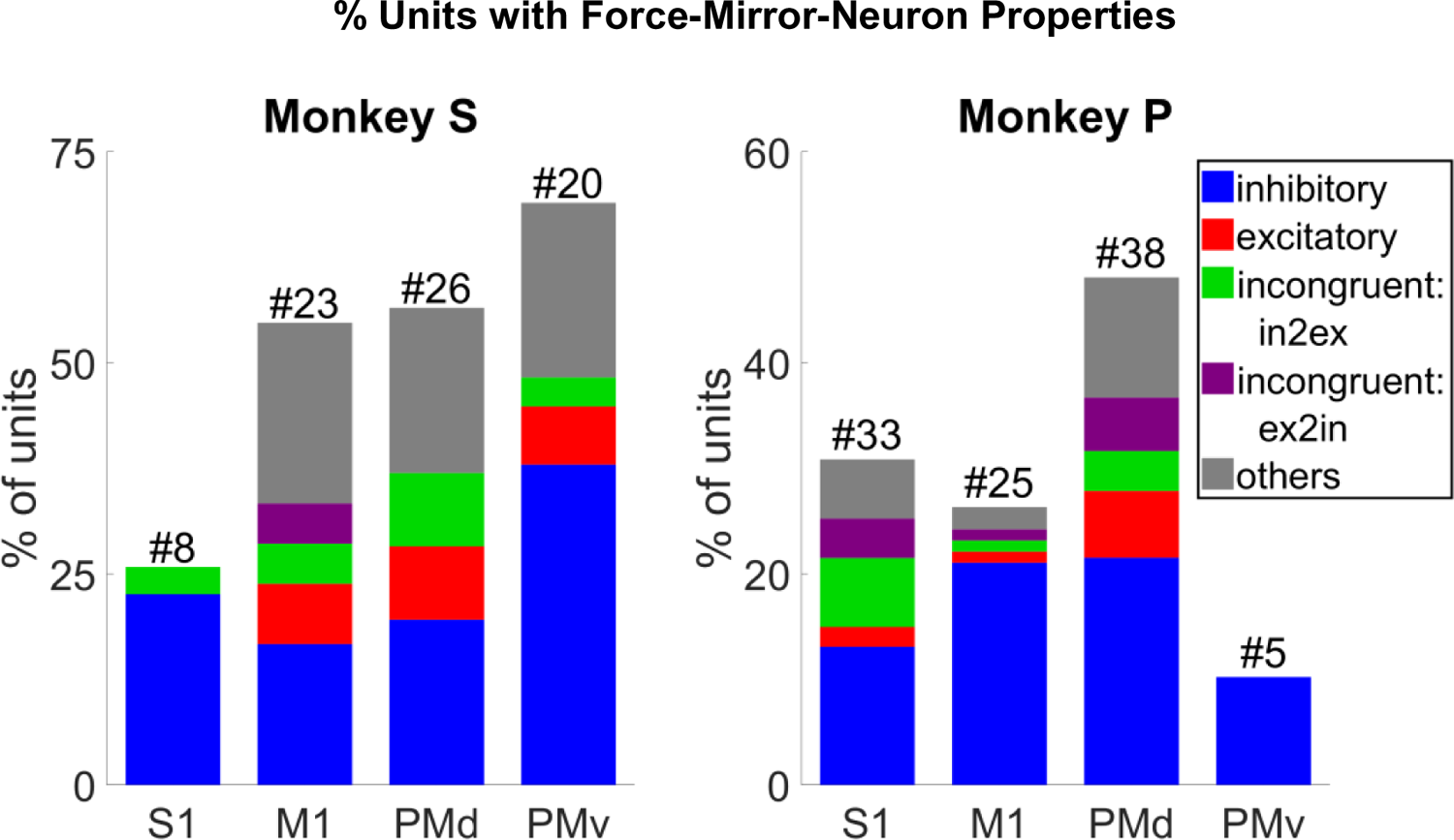
Percentage of well isolated MN single units from cS1, rM1, PMd and PMv that showed significant (F-test, BH post-hoc adjusted for number of units in the population) fit to grip-force during all four blocks of data (two manual and two observational) for each NHP. The different color bars represent congruent inhibitory (blue), excitatory (red), incongruent inhibitory to excitatory (green), incongruent excitatory to inhibitory (purple), and other (gray) unit responses. The percentage (y-axis) shows how many units (# at top) are significant for force from units present in all four data blocks. See Fig.S11-S12 for a breakdown of units by trial type.

### Distribution of Reactive vs. Predictive Mirror Neurons

The analysis in this section was conducted in order to determine if there was an obvious shift in time lag between the neural correlates of force between manual trials and observational trials. One would expect the neural data responsible for force production, or imagining force production, to lead the force output, whereas classical MN activity would lag the viewed “force” output. However, this may not be the case for predictable movements, such as used in our task, as described by others ^43^, where MNs can still lead the observed task. In Fig.7 units that were significant for force (F-test, p <0.05, eq.1) on all block types were plotted in the time bin where their correlation coefficient with force was maximum in absolute value, either negative or positive. We asked if these distributions significantly (signed Rank test, p <0.01) deviated from zero between the manual and the observational versions of both the cued and uncued tasks. There were no significant shifts in the histograms for NHP P. NHP S showed one task with significant shifts for PMd during the cued tasks. For each plot the zero-time shift shows the probability of a unit not shifting in time between manual and observational. The positive differences are the classical MN response where observational neural activity lags the visual information, whereas the negative time shifts represent the probability of the observational neural responses being earlier, or more “predictive” of moment, than during the manual trials. For more on correlational analysis see Fig.S13 and Fig.S14.

**Figure 7:**
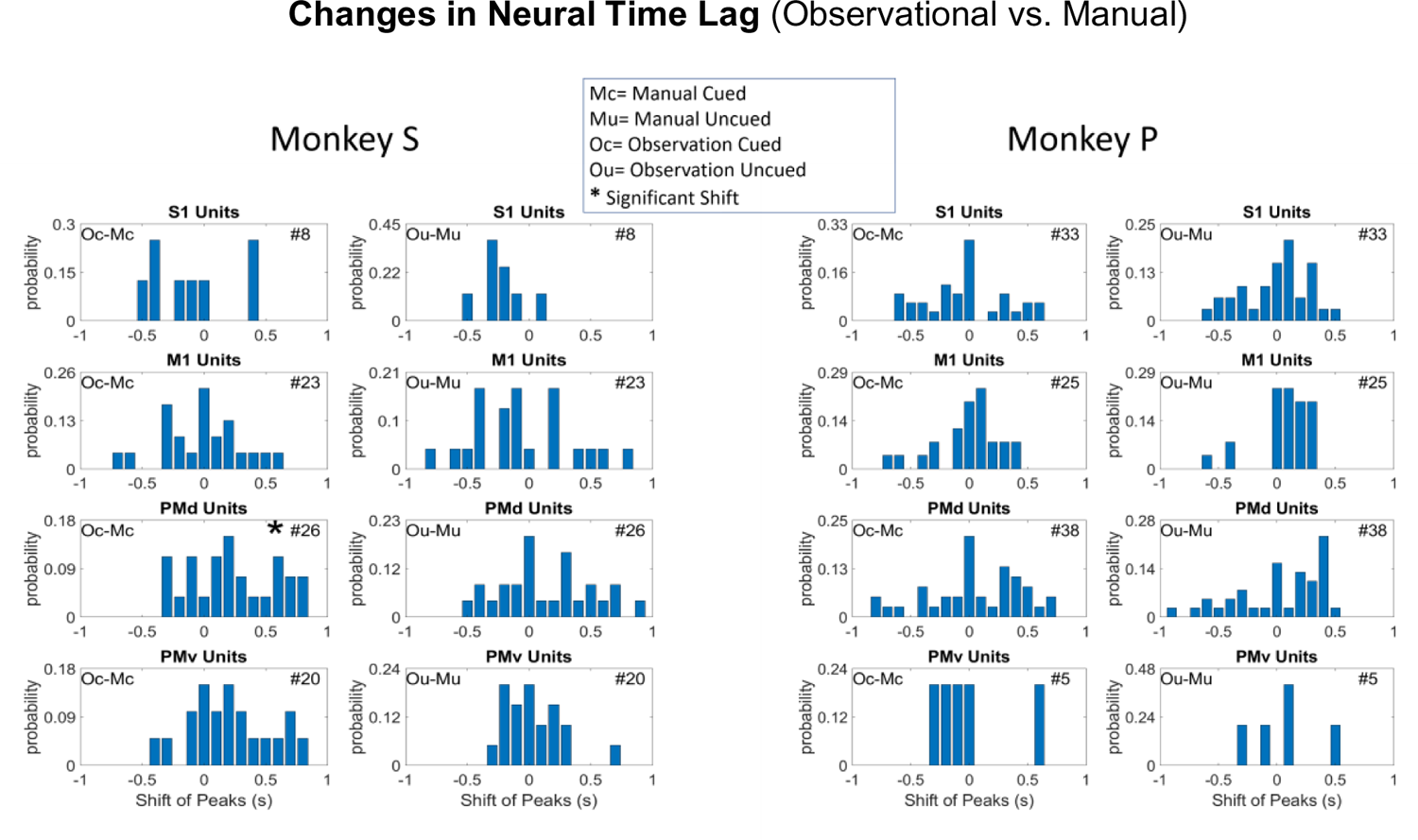
Changes in neural time lags where neural data best correlate with peak grip-force. Bar plots showing the shift of peak correlation between similar (cued/Uncued) Manual and Observational blocks for cS1, rM1, PMd and PMv cortices (cortex is labelled on the title of each subplot). The shift from a block type to another block type (manual-to-observational) is at the top right of each subplot for each NHP (Mc= Manual Cued, Mu= Manual Uncued, Oc= Observation Cued, Ou= Observation Uncued). The number of units used is indicated in the upper right corner of each subplot. The left column for each NHP shows the shift between cued blocks and the right column shows the shift between uncued block. An Asterisks (*) symbol before the number of units represents a significant (p<0.01) shift in histogram from manual to observational tasks. Probability of unit shifts was calculated by subtracting the position of the peak correlation time bin for manual from the observational tasks for the same unit. For more on this correlational analysis see Fig.S13 and Fig.S14.

### Force and Reward Modulation of Single Mirror Neurons

Figure 8 shows that a good percentage of units show MN activity encoding force during both manual and observational trials, Fig.8.a. We also show the percentage and number of units that were significantly correlated with force or reward separately, as well as units that were significant for both, but not simultaneously, which would be comodulation and is shown in Figs.9 - 10. Units in Fig.8 were significant under both manual and observational trials. Single unit activities were tracked across manual and observational cued and uncued blocks to determine the significance of their correlation with force under these different conditions. To clarify, a unit that was present during both cued manual and cued observational blocks was checked for significant correlation with force and/or reward and included in Fig.8 only if it was significant for both manual and observational trials towards possible discovery of MNs. The same procedure was performed for uncued manual and observational blocks separately. The plots in Fig.8 show the % of units (left y-axis) and total number of units (top of each bar) for each category that were identified from cS1, rM1, PMd and PMv recordings. The green bars are for reward level cued trials while the purple bars are for reward level uncued trials. The analysis windows used for reward were, for post-cue from 0s to 0.5s, and for post-result from 0s to 0.5s while for force significance the window was 0.5s starting at pre-force-onset to 0.5s after post-force-offset. Note these time windows were not overlapping, thus Fig. 8 is not describing results of force tuning modulation by reward, which is described in Figs.9 - 10.

**Figure 8.**
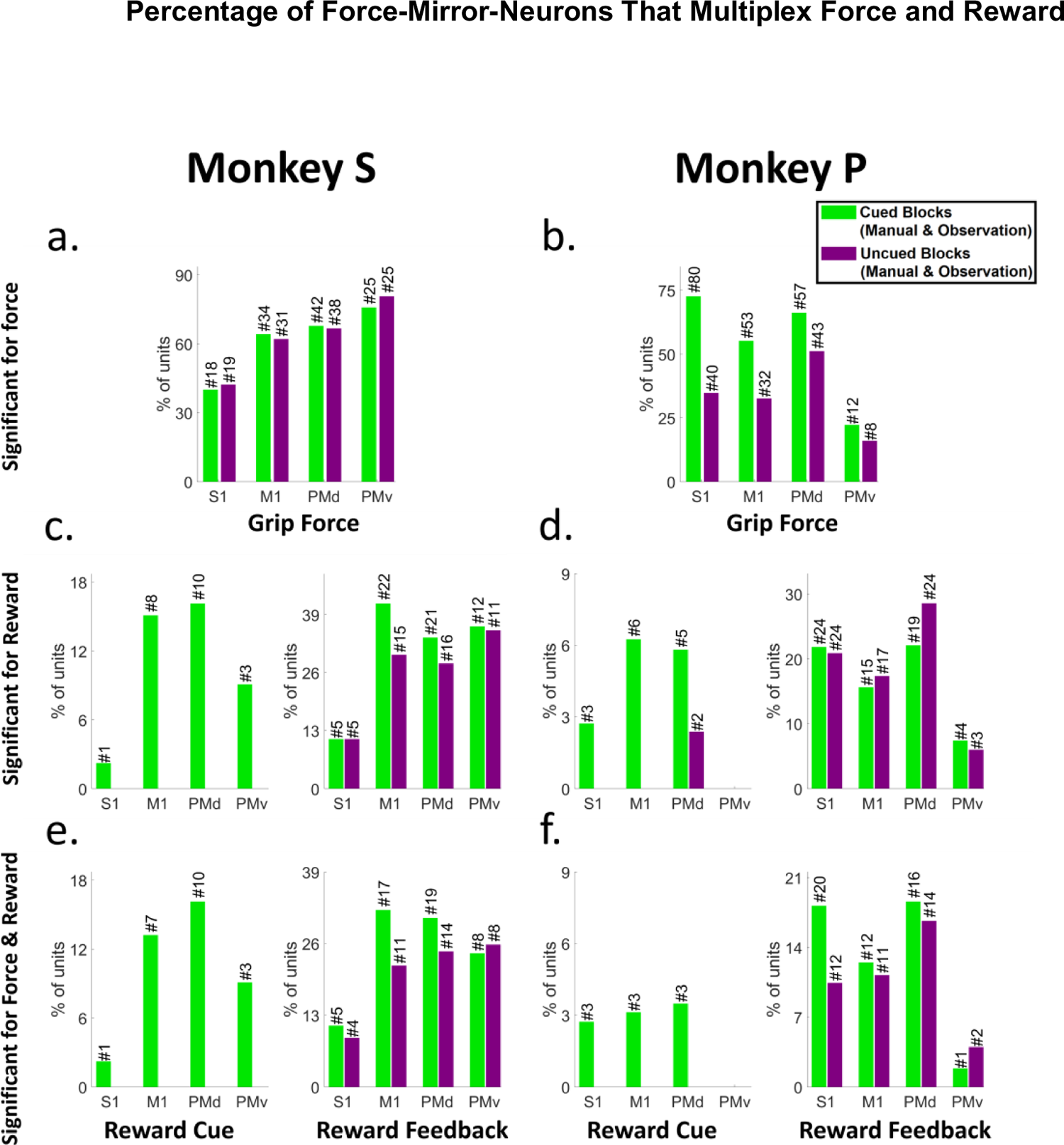
Mirror Neurons in cS1, rM1, PMd and PMv Multiplex Reward and Grip-force. The percentage of units with significant modulation via either grip-force (F test, BH method for p=0.05), reward (post-cue or post-reward, t-test, BH method for p=0.05), or both, during manual and observational blocks of a similar type (e.g. cued or uncued). Number of hypothesis tests for the post hoc BH method was the number of MNs in that brain region under study, seen above each bar. Green represents units for cued blocks and Purple for uncued blocks. Each subplot shows the percentage of significant units between cued/uncued for units showing such activity during both manual and observational blocks for Force (plot a and b), Reward (plot c and d) and significant for both force and reward (plot e and f) during separate times in the task. For subplots c, d, e and f the left plot shows units for post reward cue and right plot shows units for post reward activity after reward delivery (150ms) was completed.

### Comodulation of Force Tuning Curves by Cued Reward Level

In Figs.9-10 we are asking questions about the modulation of the force tuning curves of MNs by cued reward level. Force tuning curves for R1 and R0 trials as well as significant differences between their slopes were calculated as described in the methods using MATLAB’s ANCOVA function. F-statistics were conducted on the reward group*force interaction which expresses the difference in slopes and the p-values for those interactions were collected. Fig.9 shows the force tuning curves for example units recorded from reward cued observational blocks for each brain region (see Fig.S15 for the manual version). The relationship between R1 and R0 trials as spike rate varied with force can be seen in Fig.9. Note this is in comparison to the previous sections when force decoding meant the neural activity could be used to determine the force level, whereas here we are looking at the change in firing rate as a change in force level and its modulation by reward (encoding). The smoothed spike rate against force is shown in the left subplots while the line plots to the right show force tuning curves obtained from the ANCOVA. The two example units from each NHP, S and P, had significant differences between R0 and R1, in agreement with our previous rM1 results for manual trials ^8^, again, results in Fig.9 are for observational trials only and manual versions can be seen in Fig.S15.

**Figure 9:**
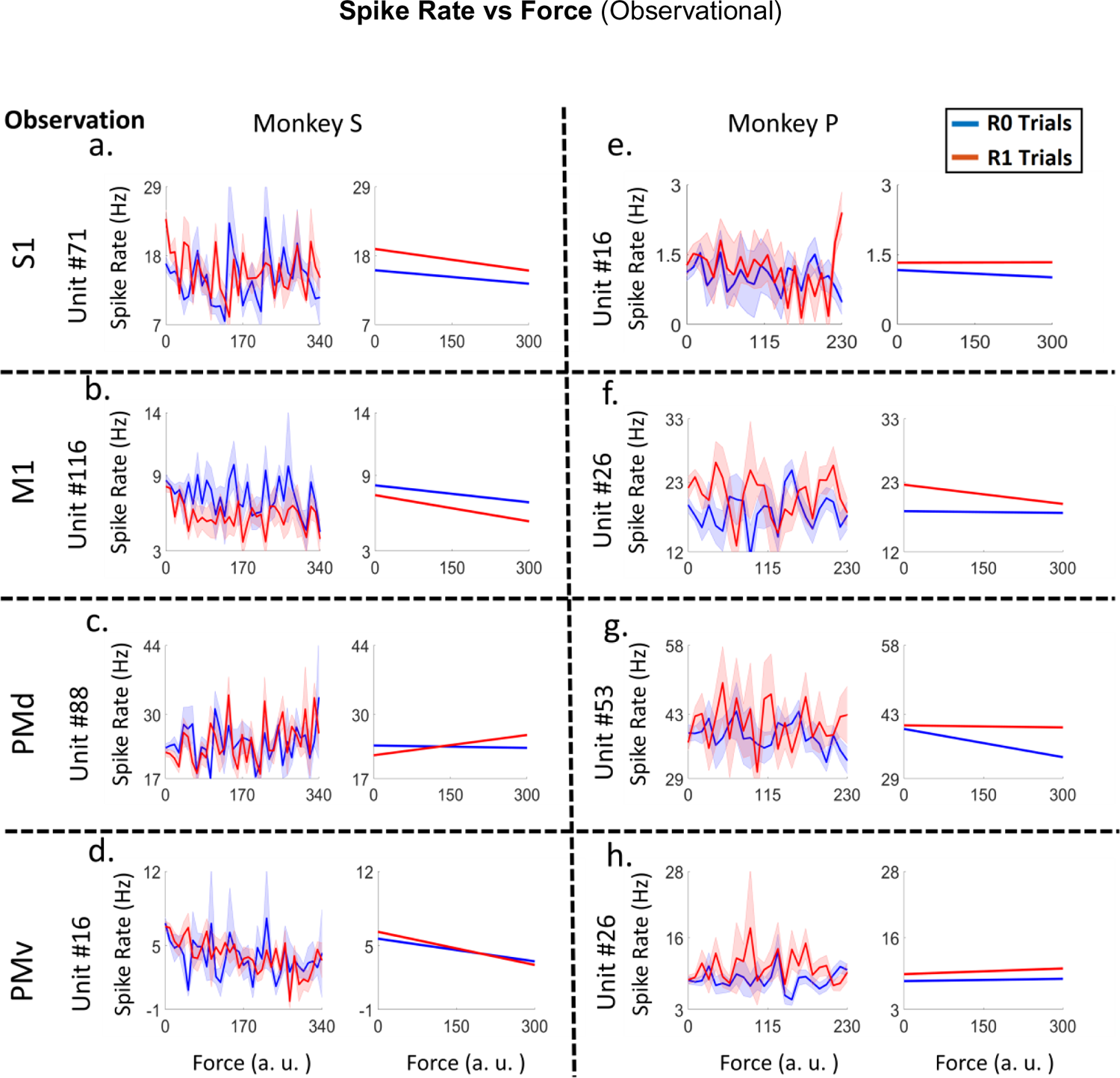
Plots of spike rate vs force (left subplots) and their linear tuning curves (right subplots) for example units from cS1 (plot *a, e*), rM1 (plot *b, f*), PMd (plot *c, g*) and PMv (plot *d, h*) cortices of both NHPs (for NHP S plots *a, b, c* and *d* and for NHP P plots *e, f, g* and *h*). The units presented had significant differences between R0 and R1 groups (ANCOVA, F-test, p<0.05) force tuning curves during observational blocks. Red lines indicate rewarding trials (R1) and blue indicate non-rewarding trials (R0). See Fig.10 for population results and Fig.S15 for the manual version of this figure.

In Fig. 10 we show results indicating the degree to which cS1, rM1, PMd and PMv not only multiplex information on both force and reward expectation during both manual and observational tasks, but how reward expectation modulates the force tuning functions that is the degree of comodulation. Fig. 10 green bar plots indicate the percentage of single units that have significant grip-force tuning curves (ANVOCA, p<0.01) for the NHP and task type indicated at the subplot title. Purple bars show the subpopulation of these units (green bars) that pass a rather stringent post-hoc false discovery rate correction (BH, p<0.01, # of hypothesis corrected for is the number of units being considered for tuning curve testing in that brain region). Red bars indicate the subpopulation of the grip-force units that are modulated for both task types (manual and observational) that is MNs. Likewise, yellow bars indicate the MN subpopulation after the post-hoc test (BH, p<0.01, # of units). Fig. 10 indicates that Comodulation of force by reward is most likely significant in manual tasks but support for the MN activity during observation of this comodulation of force by reward is rather weak (1-2% max), which could in part be due to the lower number of trials recorded during observation compounded by the lower firing rate. See tables S3-S4 for more information on the ANCOVA results.

**Figure 10:**
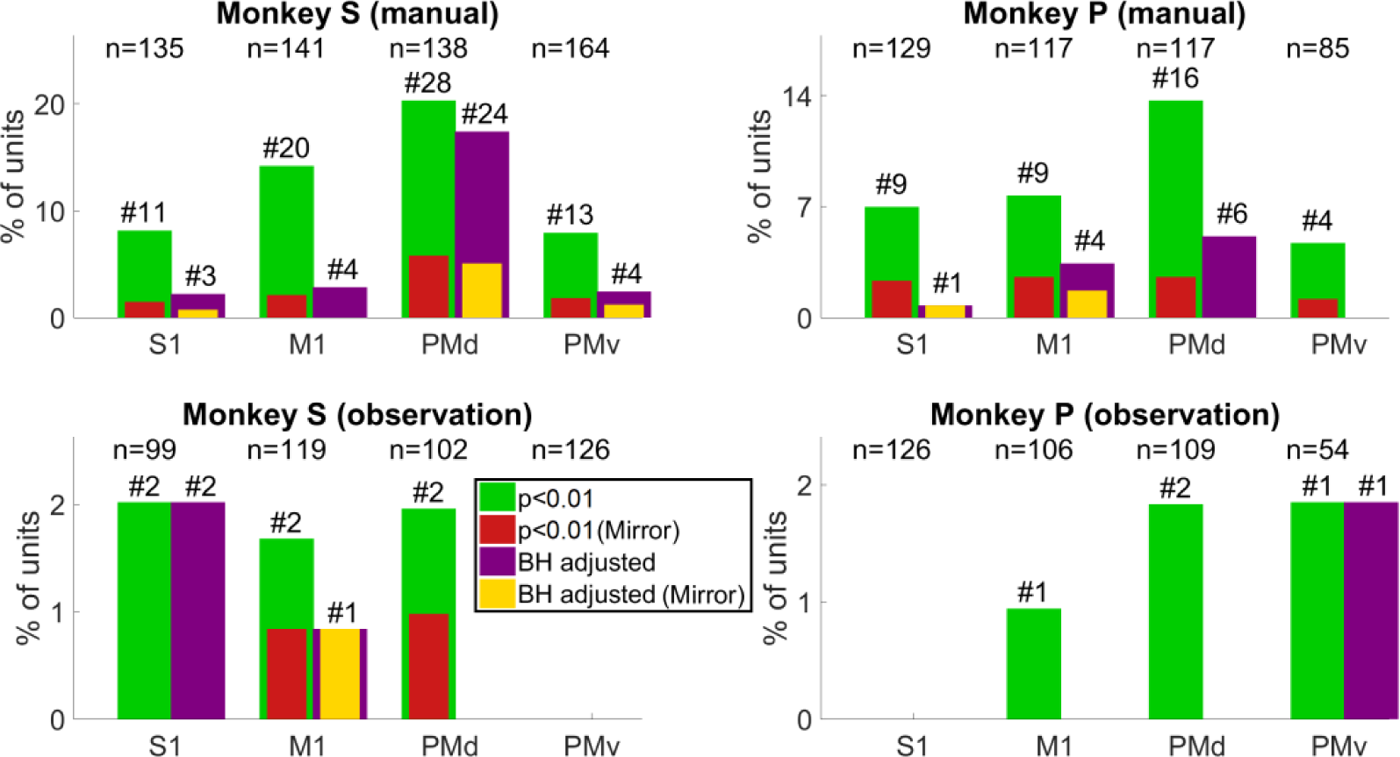
The percentage of single units with force tuning curves that are significantly modulated by reward level. The titles represent which NHP and block type are plotted, p-values were adjusted using the Benjamini-Hochberg method (BH method) correcting for false discovery rate where the correction was applied using p=0.01 and the number of hypothesis tests applied was the number of units in the population under study. The y-axis is the percentage of units and x-axis represents the cortical region. The green bar is showing the number of significant units (p<0.01, ANCOVA) and red is showing the MN population among them. The purple bar is showing the significant units after BH method was applied. The yellow bar inside the purple is showing the percentage of putative MNs after BH method. The number after the ‘#’ sign on top of each bar is showing the number of significant units for that given case and the number presented with ‘n’ is the total units recorded for that case.

## Discussion

The main outcome of this work is the clear support showing that MN’s can encode grip force (Fig.5-6 and Tables.S1 and S2), a non-kinematic variable in cS1 that is areas 1 and 2 as well as in rM1, PMd and PMv (see Fig.2). Secondly, that the MN activity during observational trials, as compared to manual trials, can shift their temporal relationship predicting/responding to visually cued force about equally in either the predictive or responsive time lag directions seen in Fig.7. Thirdly, we have shown that the firing rate in each of these regions is also modulated by reward level expectation during the post-reward-cue period for reward-level-cued trials, and during the post-feedback time period for both cued and uncued reward level trials seen in Fig.8. Fourthly, that MNs activity of visually cued grip-force can coexist (multiplex) within single units that also code for reward level seen in Fig.8. Finally, we have shown evidence, albeit weak (∼3-4% of MNs for manual and ∼1-2% of MNs for observational tasks), that the neural grip-force tuning functions can be modulated by reward expectation (comodulation) in the brain regions under study (cS1, rM1, PMd and PMv) during both manual and observational trials Figs.9-10.

To the best of our knowledge, we are reporting the first evidence of mirror neural responses to expected, or visually cued effort (grip-force), within these sensorimotor regions at the single unit level. As the NHPs had a great deal of experience with the cued grip-force task we expect the mirror responses were due to the visual cueing of the force targets that the NHPs had come to understand based on their manual trials. In Fig.7 we showed that the shift in peak correlation between cued force and neural time lag could be in either the direction expected for motor production, or that expected from sensory stimulation/gating, which would be more in line with the initial MN work ^1^, and the former more in line with movement rehearsal ^2^. We predict that such force MN activity could be seen in more natural settings if the context was understood by the NHPs, such as lifting heavy vs. light objects that they were familiar with, or perhaps when the NHP squeezes a deformable object, or observes such a grasp, again with knowledge of the object’s physical properties. However, further work is needed to show such activity exists under more natural conditions within each of these regions. It should be noted that this type of prior knowledge being necessary for MN activity patterns is not new. Mirror neurons that encode subjective value in addition to grasping information clearly need the NHP to have familiarity with the object in some way to have formed a subjective value ^46^.

We have shown that reward modulation can occur post-cue, when the cue indicates the reward value of the current trial, as well as post-feedback when the reward level that was cued is delivered. When there is no reward cue given in uncued blocks there is no “post-cue” reward modulation as expected, but there is still post-feedback modulation in each of the 4 brain regions, and again during both manual and observational trial types. Based on the evidence we have presented, we see that cS1, rM1, PMd and PMv contain units that encode reward expectation and reward itself, as has been shown previously for the task used in this work as well as other tasks for rM1 ^6, 8, 23, 28^, indicating this reward modulation may be a generalized broadcast signal to regions involved in the ongoing sensorimotor planning and production of movement, as well as during observation of such movements, in a manner that can be predictive, as expected during mental simulation ^2^, and responsive to the visual stimuli, as expected for classical MNs (Rizzolatti et al. 1996; Gallese et al. 1996). In addition, we have now shown such reward modulation simultaneously for rM1, cS1, PMd and PMv during both manual and observational trial types for the same single units tracked between tasks.

There are several limitations to the work presented here that should be addressed in future work. EMGs were not available for these particular datasets that allowed us to track single units between manual and observational trials, and therefore we can’t state with certainty that the NHPs were not activating their arm muscles covertly, however, they were not making obvious movements, and did not have access to the force transducing handle, in addition, our previous studies found no correlation between EMGs and curser motion in observational trials of a reaching task ^28^. In addition, we show in Fig.S16 for both NHPs there was no significant EMG activity during observational trials on different days when the EMG signals were usable. The grip-force output in the current task was generally phasic in nature with a bell-shaped profile, even if the cued amplitudes were different, the force profiles were still fairly stereotyped, and so it is possible that some other phasic neural response was allowing the regression models to predict force output, which was bell-shaped, by some non-force related phasic response, such as that related to the cued reward information. However, the regression models did not fall apart in the uncued reward level task, which indicates at least cued reward does not explain our force decoding results. In addition, when we conducted analysis on the time series of just the peak force amplitude of each trial, we still obtained positive results showing MNs for force, although clearly not as strong as when using the fuller dataset including the force trajectories (see Fig.S19). Additionally, in Fig.S20 we show significance holds for several measures on the manual data between R0 and R1 trials post data pruning to keep the trials between these two categories statistically indistinguishable.

Previous work has suggested that MN activity is not seen in S1 ^47^, however they did see non-frequency discriminative activation of S1 due to auditory stimuli. Others have suggested that S1 is in fact modulated in a MN manner, however, they utilized fMRI ^48^. Histological studies where MN brain regions were injected with tracers did show that IPL injections led to staining of S1 ^49^, and these studies by Meyer et al. and Bruni et al. would imply our results are perhaps expected. The data we present for MNs encoding grip force are very clear and significant as are the results showing that these MNs multiplex cued reward level, or reward expectation and force, however, the results for comodulation of force tuning curves by reward were less convincing, but still significant.

The work presented here has a practical application past basic neuroscience. We believe it is important to continue gaining a better understanding of encoded information, such as reward, within the sensorimotor cortices toward the development of a close-loop brain machine interface (BMI) for the restoration of motor control and beyond, such as towards a better understanding and tracking of psychological state of the individual. BMI neural signals are often recorded and decoded from a subset of cS1, rM1, PMd, and PMv while sensory feedback is obtained either by natural vision, or stimuli routed to cS1, such as via the thalamus ^50^ for somatosensory feedback. It has recently been shown that reward expectation can change directional and force based tuning curves in rM1 and cS1 ^8, 23^, and here we have shown that all of these brain regions are influenced by reward expectation, effort, or sensory feedback on expected effort, and therefore research into these relationship is warranted. Differentiating between mirror activity in these regions and intentional activity for movement is key towards making BMIs more stable with respect to the users intended movements compared to observed movements, and our future work will address these issues.

## SUPPLEMENTARY INFORMATION

### Supplementary Information

#### The number of single units recorded for each cortex

The figure below shows the total number of single units recorded in all the cortices used (S1, M1, PMd, and PMv) and for each of the block types (MC, MU, OU, and OC). The block types are color coded as seen in the legend in the right subplot. The left subplot shows the number of units from NHP S and the right shows from NHP P. The x-axis is the cortical region and the y-axis is the number of units.

#### Decoded Force R-square value with and without smoothening (Fig.S3)

We applied MLR (Multiple linear regression) to decoded grip force from smoothened spike rates (shown in main text figure 5). In order to show that the smoothening was not playing a role in our decoding results we also tested the square root transformed spike rate without smoothing to decode the force following the same procedure described in the method section. The following figure shows the R-square values of force decoding with (red) or without (blue) smoothening.

#### Grip Force prediction without Square root transformation (Fig.S4)

In the following figure S4, we show a comparison between the grip decoding accuracy with R-square values between decoded grip force on the test set of data and the actual grip force applied (manual) or observed (observational). The bar plots showed that the R-squared doesn’t deviate much when a square root transform is applied (blue) on the spike rate from the case when the transformation was not applied (Red). The figure 8 in the main text shows that NHP S had a higher percentage of significant units, especially in M1, PMd and PMv cortex due to reward cue compared to NHP P.

#### Algorithm for Linear Regression

The following algorithm shows the two-step process of collecting a subset of units significant for force to achieve a comparatively better prediction of force using linear regression. **Fig.S5**

#### The Force duration for each data block

The durations of applied grip force in R0 (blue) and R1 (red) trials are given on the following box plots for each of the block types (MC= Manual Cued, MU= Manual Uncued, OC= Observation Cued and OU=Observation Uncued). The box plots are showing the highest (upper line), 75^th^ percentile, median, 25^th^ percentile and the lowest values for force duration of all the trials. A paired t-test (p-value <0.05) was performed to check if the R0 peak force was significantly different from R1 trials. The R0 and R1 force durations are significantly different only for the case of MC block of Monkey S.

The y-axis is representing the force duration in seconds. The black line and asterisk (*) on the top are showing the significant difference between R0 (blue) and R1 (red) trials.

#### Peak grip force for each data block

The applied/observed peak grip force on each manual/observational blocks are plotted on the following figures. The Red is representing the R1 trials and blue is R0 trials. The block type (MC, MU, OC, OU) is mentioned in the x-axis. A paired t-test (p-value <0.05) was performed to check if the R0 peak force was significantly different from R1 trials. The significance test showed no p-value less than 0.05 and hence no asterisk (*) sign was plotted in any of the pairs. The minimum threshold of force for monkey S was 150 and P was 100 for the trial that had lowest force peak threshold. If the value goes below that value, the trial will be failed. The value mentioned is the minimum value for different trails a random value was added with the to increase the minimum threshold from 100 (monkey P) and 150 (monkey S).

#### Reaction time to apply grip force

The reaction time to apply grip force for R0 and R1 trials is plotted on the following subplots. The manual blocks for Monkey S (left) and Monkey P (right) were plotted. The blue and red color represents R0 (blue) and R1 (red) trials from the same data block (MC, MU). The asterisk (*) on the top of the black line indicates the reaction time is significantly different between R0 and R1 trials (paired t-test, p-value <0.05). The reaction time in this case is the difference between the times, NHP was allowed to apply grip force (start of the grip force scene) to the time when they applied 10% of the peak grip force. From the following plot the reaction time is significantly different only for the manual cued block for Monkey S.

#### The number of trials in each data block

The following figure shows the number of trials for each data block that were used for analysis.

#### Observation Results on force (Fig.S10)

This figure shows raster plots of some more example units from each of the 4 cortices during grip force onset and offset taken from observation blocks showing the full average force time period with both increases and decreases to force onset.

#### Responses of Mirror neuron activity due to grip force for different data blocks

The following figures are showing the number of MN units in terms of their activity during different blocks. These units are already shown in figure 6 of the main manuscript but the other category was not elaborated there, which is given in detail in figure S11 and S12. For analysis, four data blocks for each NHP (NHP S and NHP P) were used. The abbreviations for the block types are, MC=Manual Cued, MU=Manual Uncued, OU=Observational Uncued and OC=Observational Cued. For figure 6, we used the peak value from the absolute correlation coefficients between grip force and each of the 10 bins of spike rate (100ms from −0.5ms pre grip force spike rate to 0.5s post, described “Grip force trajectory prediction” section inside method) to detect inhibitory or excitatory activity. We considered a unit’s representation excitatory if the actual value of that peak absolute correlation was positive otherwise it was considered inhibitory. We detected congruent units among the significant units that showed similar activity in all four data blocks of either inhibitory (blue) or excitatory (red) spike rate in relation to grip-force. Incongruent units showed opposing behavior between manual and observational tasks, such as excitatory during manual and inhibitory during observational, purple, or inhibitory during manual and excitatory during observational (green). Other (gray) units didn’t follow any simple pattern during all four data blocks.

In the following two figures (S11 and S12) we are showing the “other” units in detail for each of the combination pattern. On the x axis label is showing four numbers for four data blocks in the order of (MC-MU-OU-OC) and a number ‘0’ represents inhibitory activity for that block and ‘1’ represents excitatory activity. So, for example the bar with the first label ‘0000’, shows the number of units that showed inhibitory spike rate during grip force activity during all four data blocks (MC, Mu, OU and OC). The bar colors are kept as like figure 6 and given as the legend. The bars that contain the other unit category from figure 6 are kept gray colored here.

#### Time Dependent Correlation Analysis (Fig.S13)

The time dependent peak correlation plots for all the blocks are given below. Only units significant for force encoding that were tracked through all manual and observational blocks are shown below.

#### Distribution of Significant Correlation Time Lags Between Neural Activity and Force (Fig.S14)

The following figure is showing the time shift of significant peak correlation from manual cued blocks to manual uncued blocks and from observation cued blocks to observation uncued blocks. See Fig.7 for more of such plots between manual and observational.

#### Regression on additional observational blocks

table S1 shows the regression data analysis for NHP S and P. The table contains the R-square value for model fit given in equation 1 and the corresponding p-value and F-statistic. The R-squared value for prediction and its p-value is also shown on the table. This table contains two additional observational blocks that were recorded on a different day from the data shown in the main text.

#### EMG Analysis

Thee data used in the main text had corrupted EMG and we were unable to confirm whether the NHPs were still using their arm muscles during the observation task, however, from observation notes of the experiments, and the clear decrease in activity seen in Figs 3-4 of the main text, and the fact that the NHPs could not reach for, or touch the force transducing handle, we do not believe the NHPs were physically rehearsing grip movements during observation. In addition, as seen below in Fig.S16, EMG on other days from these same NHPs was available during observation trials and did show any obvious activation patterns in the mean activity, however, it appears there was some small and insignificant increase in biceps for NHP P. The neural results of these below data sets held to that shown in the main text and figs. This does not rule out the NHPs covertly imagining movement, which was addressed in the main text.

#### Units significant for Extensor Muscle EMG

Figure S13 shows, most recorded units are showing negative correlation with grip force and we speculated that it could be a reason of they are connected to the extensor muscle group in the forearm. To investigate further we wanted to observe units that are showing negative correlation with force has any positive correlation with the EMG recorded from extensor muscle group. We used manual data block recorded on another day for monkey S which had EMG recorded for the extensor muscle from forearm. Unfortunately monkey P extensor EMG was not recorded on the other days. For correlation EMG and spike data was taken from all trials and concatenated. We used binned (100ms) spike data taken from 500ms pre force onset to 1000ms post and measured the spearman rank correlation with corresponding binned envelope of the EMG data from extensor muscle of forearm. The units that showed significance for force fit and a negative correlation with the grip force were further investigated to observe their correlation with extensor muscle EMG with the procedure described above. The percentage of such units that has positive correlation with extensor muscle EMG and negative correlation with grip force for NHP S is shown in figure S17. Figure S18 is showing the mean EMG and spike rate for an example unit with the raster plot.

#### Significant units for Peak Grip force

As force profiles were stereotypical smooth Gaussian-like profiles, we wanted to make sure our regression model fits and predictions were meaningful, and not solely due to a unit having a phasic response that could be used to fit the stereotypical waveform. Therefore, we determined how neural activity correlated with the peak values of the Gaussian-shaped force profiles during each trial. We considered force values around the peak force applied on each trial (three values around the peak including the peak value) and we placed neural data, centered around each of them, into ten 100ms non-overlapping bins from 500ms pre peak force to 500ms post peak force neural activity. Each MN (figure 6 on the main manuscript) was tested for peak force significance (F-test, BH corrected for number of population that is the number of corresponding units) by fitting a linear regression model with peak force values and spike rate for each of the ten bins.

The blue bars are showing the units with Force-Mirror-neuron properties (Figure 6 inside main manuscript). The red bars on figure S11 are showing significant units for peak grip force activity among the force-mirror-neuron units.

#### Behavioral Summary

We used two NHP for our experiment, one male *Macaca* Radiata (NHP S) and one female *Macaca* Mulatta (NHP P). Both NHP performed manual and observational task. During the observational version of the task a plexiglass box restrained their hand from reaching the grip force sensor. For the manual version of the task they were able to reach the force sensor. During manual trials when they were cued to apply grip force, they reached to the grip force sensor and at the end of a trial they usually rested their arm on the base of the grip force sensor. NHP S achieved success rates of 77% (cued manual) and 82% (uncued manual) during the two manual-task blocks, while NHP P achieved rates of 58% (cued manual) and 72% (uncued manual). This comparatively low success rate for the manual block when they were cued for reward at the beginning shows that both NHP had an idea about the reward information. Between the two monkeys, performance of S during the trial was affected more by reward cue information compared to P. Figure S6 shows the force duration is significantly different between R0 and R1 trials for NHP S which is not same for NHP P. Also, from figure S7 we can see the reaction time is significantly longer when NHP S completes R0 trials compared to R1 trials. Both results are indication that NHP S task performance was more dependent on reward compared to NHP P. One thing that should be mentioned that, for NHP S, although reaction time and grip force duration was significantly different the peak force applied on each trial was not significantly different (figure S11) since the applied grip force was supposed to be within a boundary value cued during task.

#### Mirror neuron distribution from the array center

The position of the mirror neurons with respected to the electrode they were recorded from is shown on the Fig.2 on the main manuscript. To observe any non-uniform distribution of the mirror neurons with respect to the center of the array we applied Rayleigh test which used the angular position of all the mirror neurons from the center of the array and test the hypothesis if the angles of the mirror neurons are uniformly distributed or not. For each cortex we applied the test for both NHP individually and also we combined data from both NHP for each cortex and applied the statistics again. The table S5 shows the p-value recorded from the statistical tests. Only M1 cortex showed significant for all cases and the mean direction for M1 is toward (Fig. 2, Main text) pre-motor cortex.

#### Similar motor behavior data analysis for NHP S

Figure S6 and S8 show significant differences between force duration and reaction time for R0 and R1 trials for NHP S, MC data blocks. Note no such differences were noticed for NHP P. We wanted to make sure that even if we only used the trials that have similar motor behavior during R0 and R1 trials from MC block, the neural activity would still show significant reward related differences for individual units. We pruned trials from MC data block (NHP S) that had higher force duration (>2s) and reaction times (>1.5s) and the remining trials showed the difference between these pruned R0 and R1 trials were not significantly different for both duration and reaction times. See Fig.S20 figure legend for the results of this comparison.

**Figure S1:**
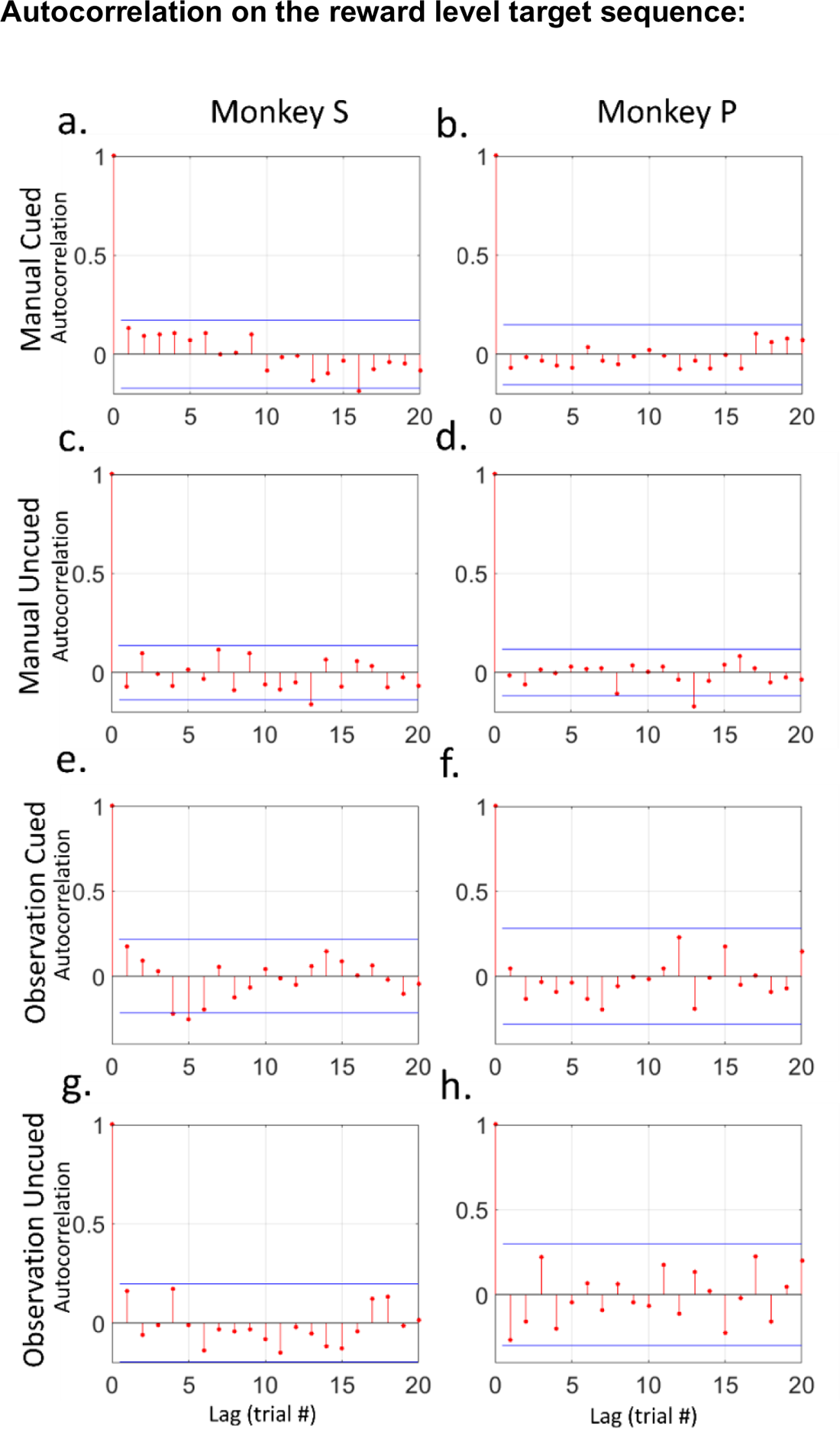
Autocorrelation on the reward level target sequence (R0 and R1) of trials for all data blocks. Plot a, c, e, and g are for NHP S and b, d, f, and h is for NHP P. The block type is given on the right side of each row. The x-axis represents lag in trial numbers and y axis represents corresponding autocorrelation.

**Figure S2:**
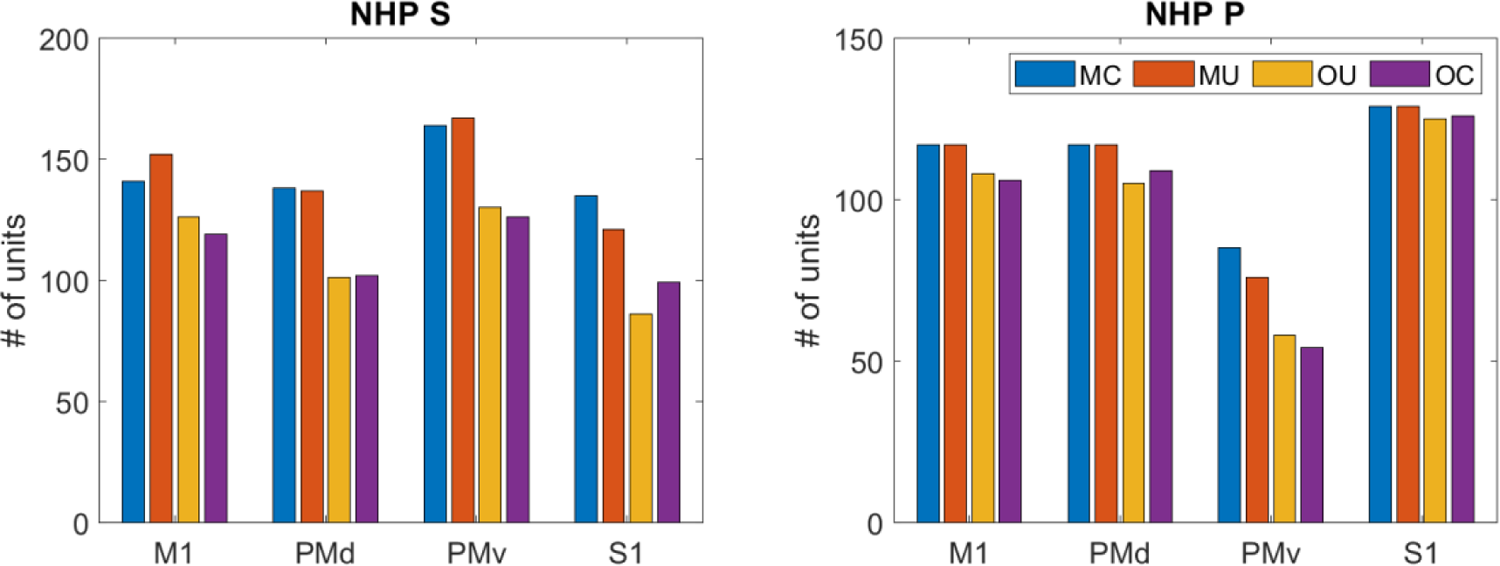
The total number of units recorded in all four cortices (S1, M1, PMd, and PMv) for both NHP S (left) and NHP P (right). For each cortex there is four columns for four data block types used (MC, MU, OU, and OC) that are color cued as in the legend on the right subplot. The y-axis represents the number of units.

**Figure S3:**
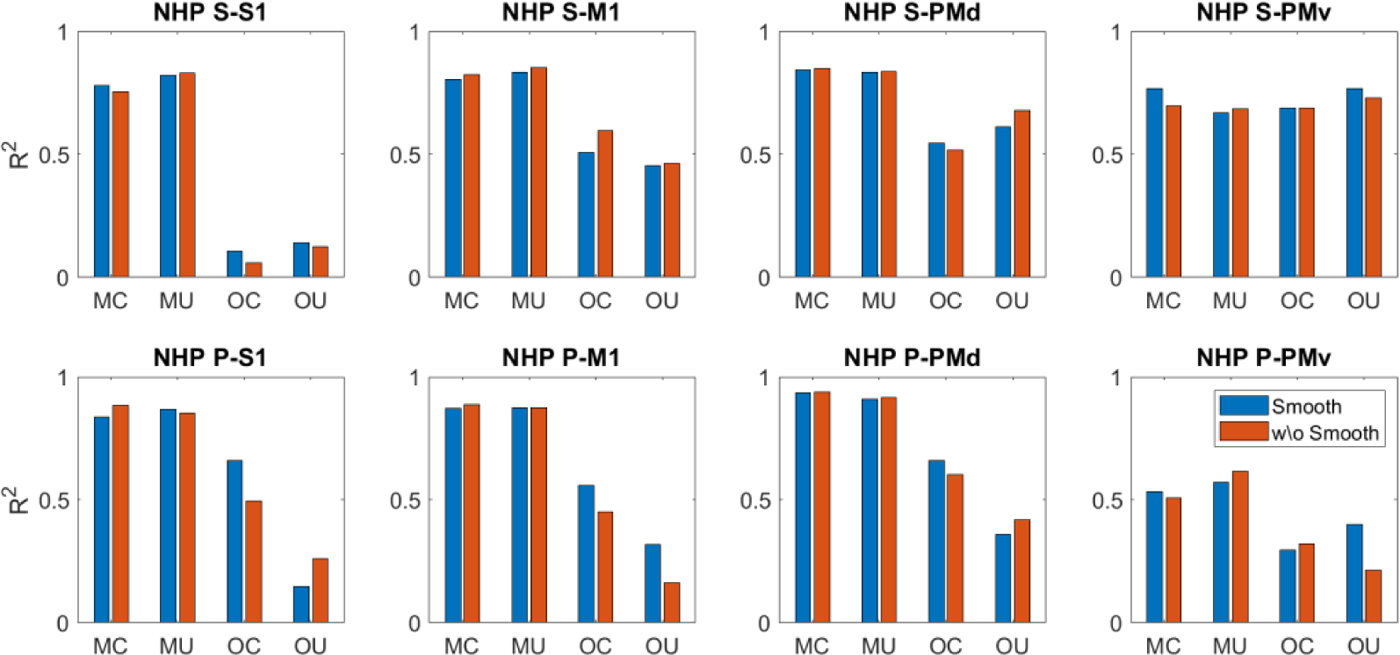
The R-square of the decoded grip force with (blue) or without (red) smoothening was applied on the spike rate data. The rows of subplot are showing data from S1, M1, PMd and PMv cortices of NHP S and bottom row is same for NHP P.

**Figure S4:**
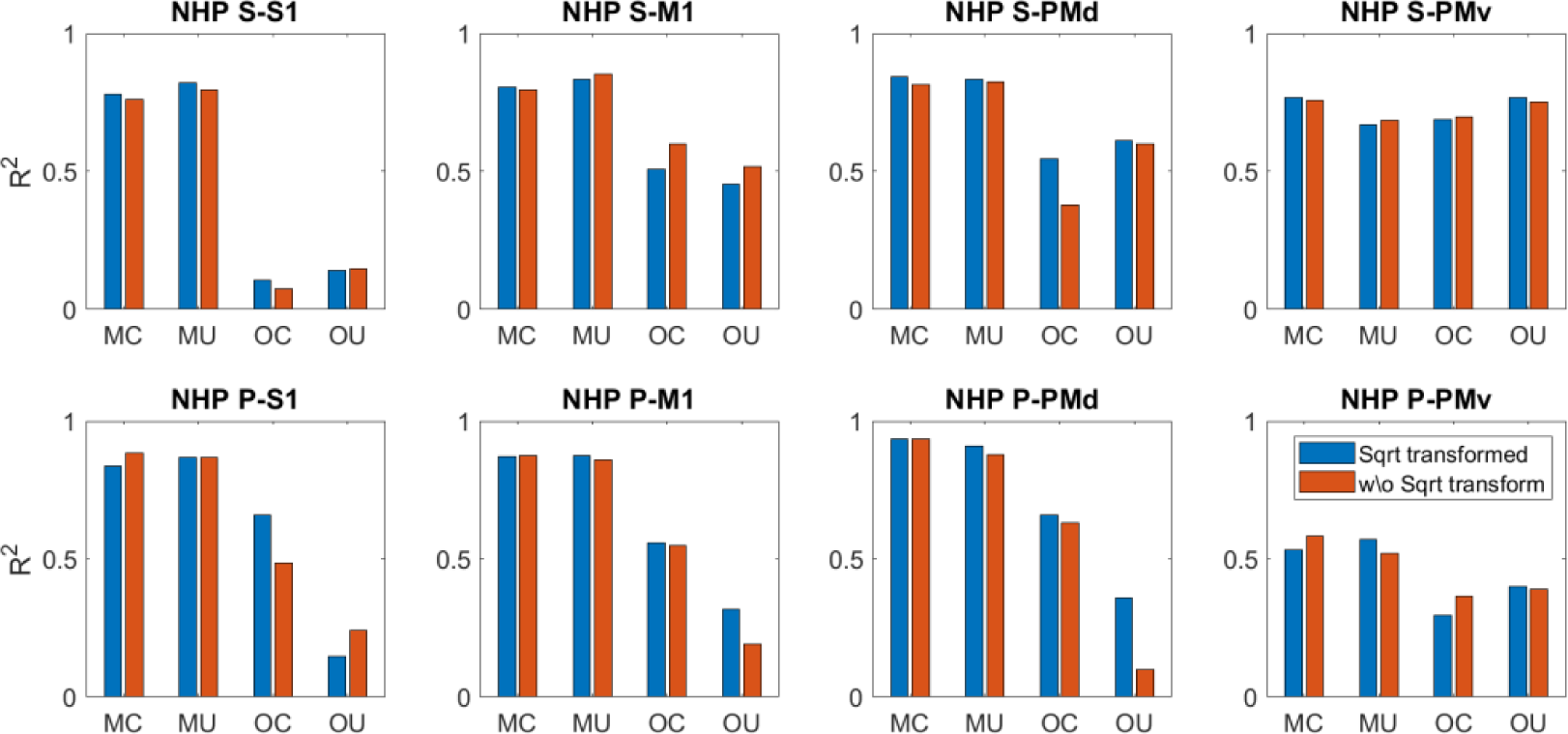
The R-square of the decoded grip force with (blue) or without (red) square root transformation was applied on the spike rate data. The rows of subplot are showing data from S1, M1, PMd and PMv cortices of NHP S and bottom row is same for NHP P.

**Figure S5:**
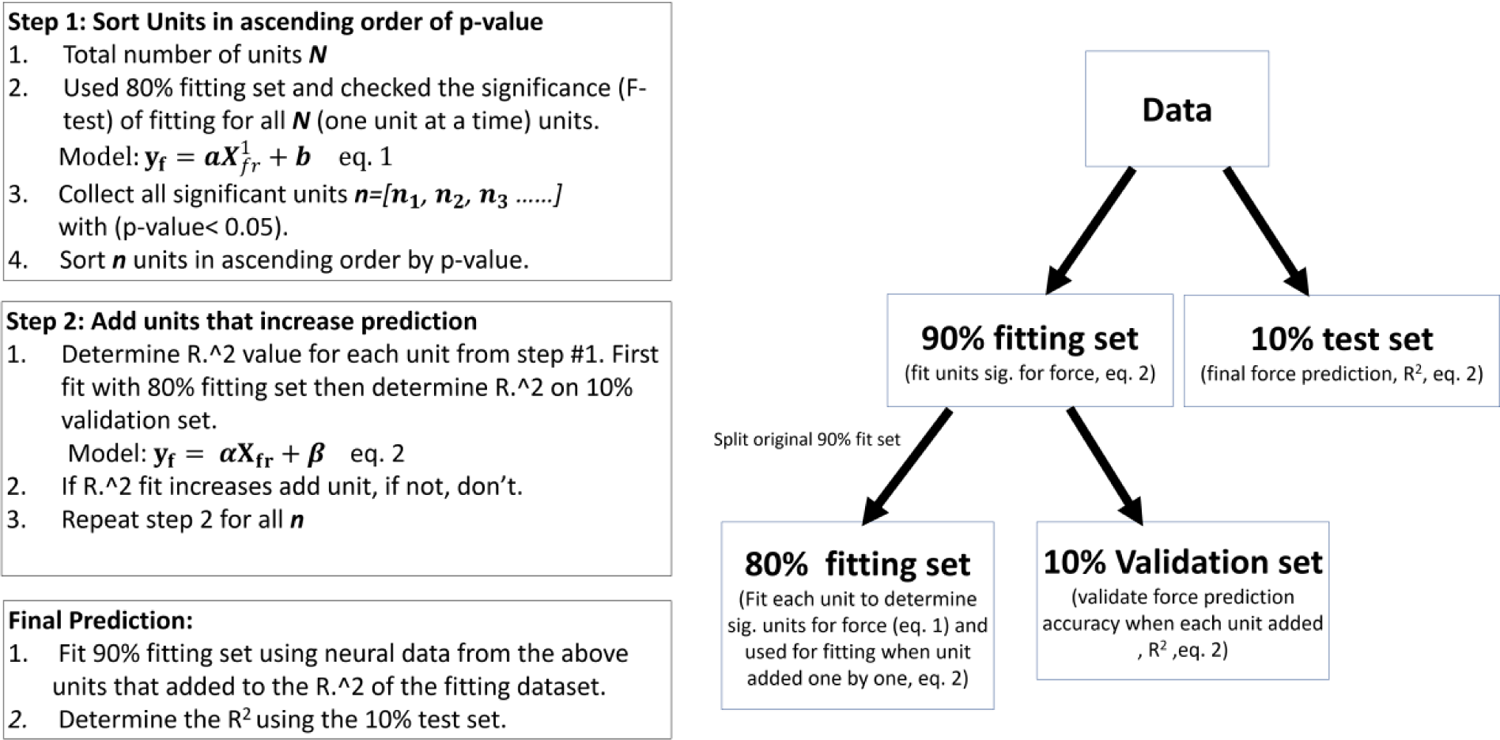
Algorithm for linear Regression

**Figure S6:**
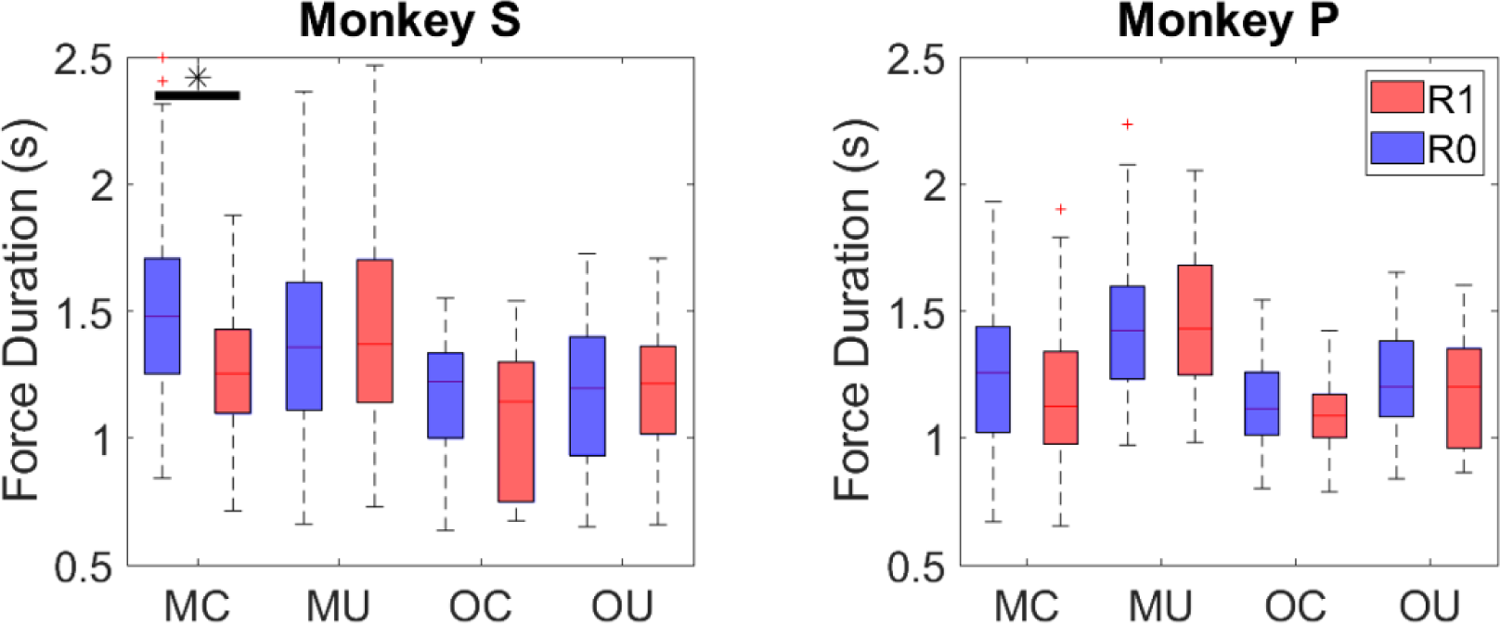
The box plot is showing the median, maximum and minimum values with 75^th^ and 25^th^ percentile of the grip force duration of all the trials on each block individually. The x-axis is showing the data block type (also shown on the legend which represents same information).

**Figure S7:**
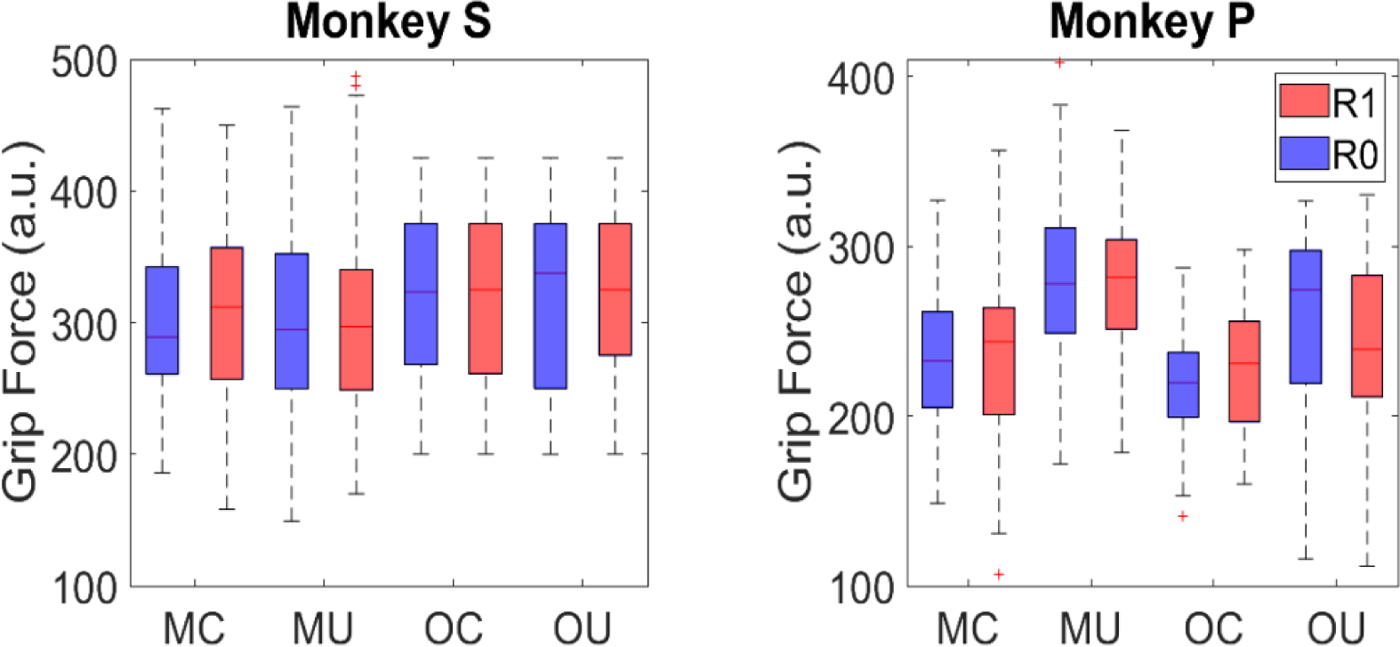
The peak force from each block type is plotted on the following box plot and for each block type the R0 (blue) and R1 (red) trials are plotted separately for NHP S (left) and NHP P (right). The x-axis is showing the block type (MC, MU, OC and OU). The y-axis is representing the value of the grip force applied with arbitrary units. The grip force values from R0 and R1 trials were significantly different for any of the case and hence no asterisk (*) is present.

**Figure S8:**
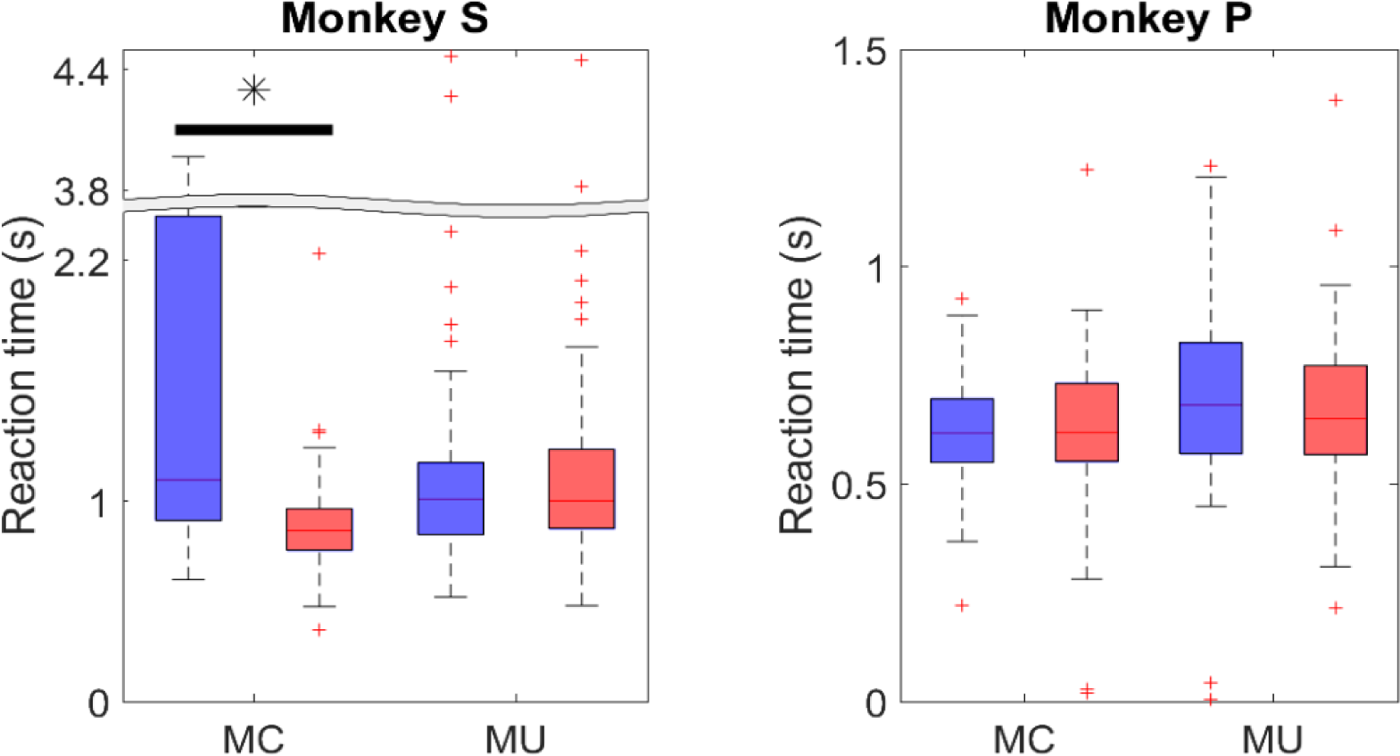
The reaction time for R0 (blue) and R1 (red) trials for all the block type is given on the following plots. The x-axis is showing the block types MC, MU for the case of manual blocks. The y-axis is showing the reaction time in seconds. The asterisk (*) on the top of each pair represents they reaction time is significantly different between R0 and R1 trials. The left subplot is for Monkey S and right subplot is for Monkey P.

**Figure S9:**
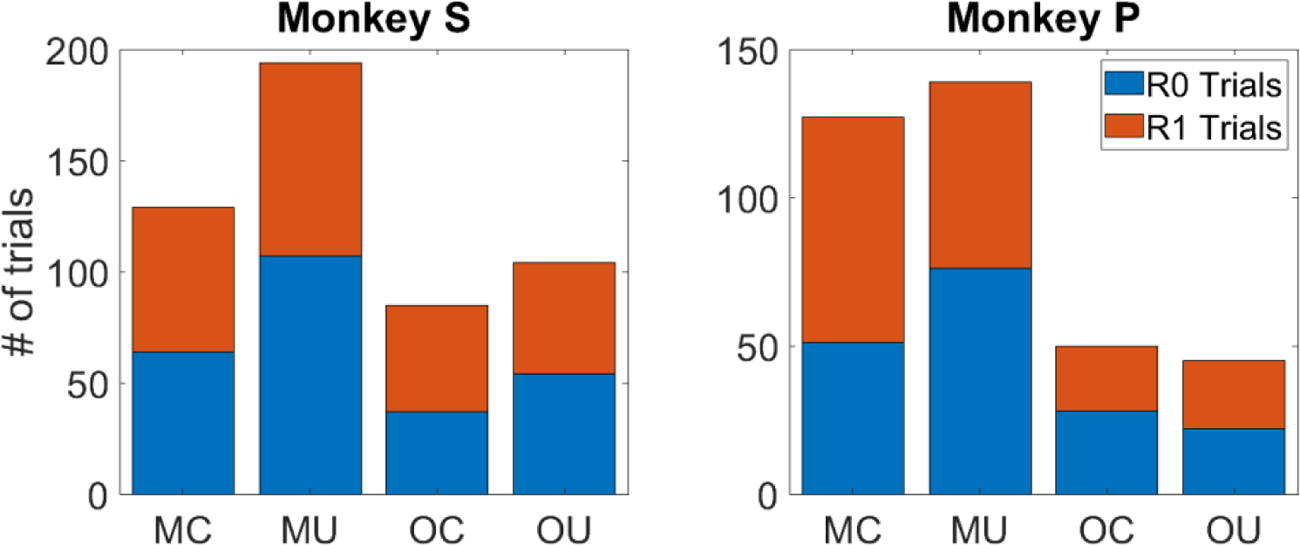
The number of trials used for the data analysis is shown on the figure above for monkey S (left) and monkey P (right). Each bar represents the total number of trials and the blue part of the bar is showing R0 trials where the red part is for R1 trials. The x-axis is representing the block types (MC, MU, OC and OU) and y-axis is showing the number of trials.

**Figure S10:**
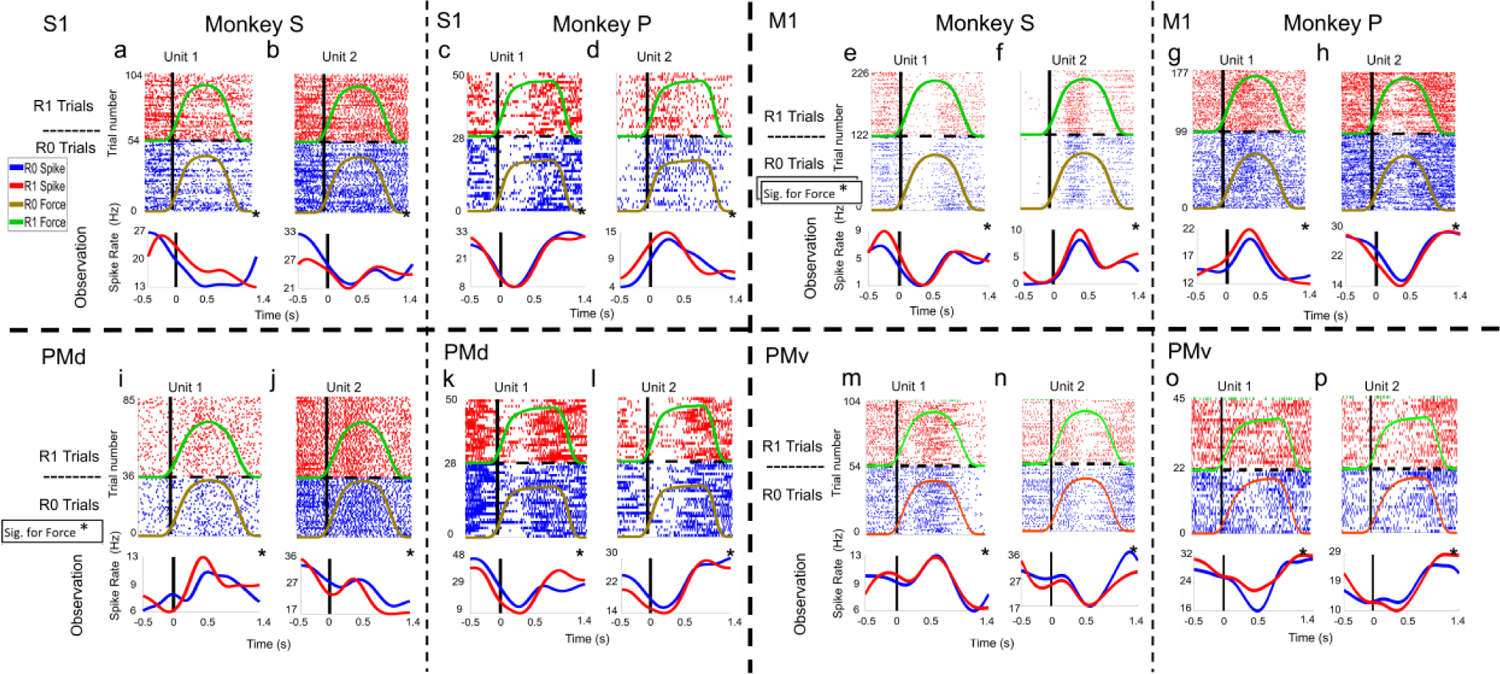
Raster plots for units from S1 (plots a, b, c and d), M1 region (plots e, f, g and h), PMd (plots i, j, k and l) and PMv (plots m, n, o and p) are shown for NHPs S and P. These plots show the grip force values expected given the cued force targets shown to the NHPs during observational blocks. The black bold line on each raster and spike rate plot shows the onset of “force” and the dotted horizontal line divides R1 (top, red) and R0 (bottom, blue) trials. On each neural data subplot, the x-axis represents time in seconds and for raster plots y-axis represents trial number (upper part of the subplots) and for the spike rate plot y-axis is spike rate in ‘Hz’ (bottom part of the subplots). The Asterisk (*) symbol on the right top corner of the mean spike rate plots represents, that unit was significant (F-test, p<0.05) for force fit using linear regression model in eq 1.

**Figure S11:**
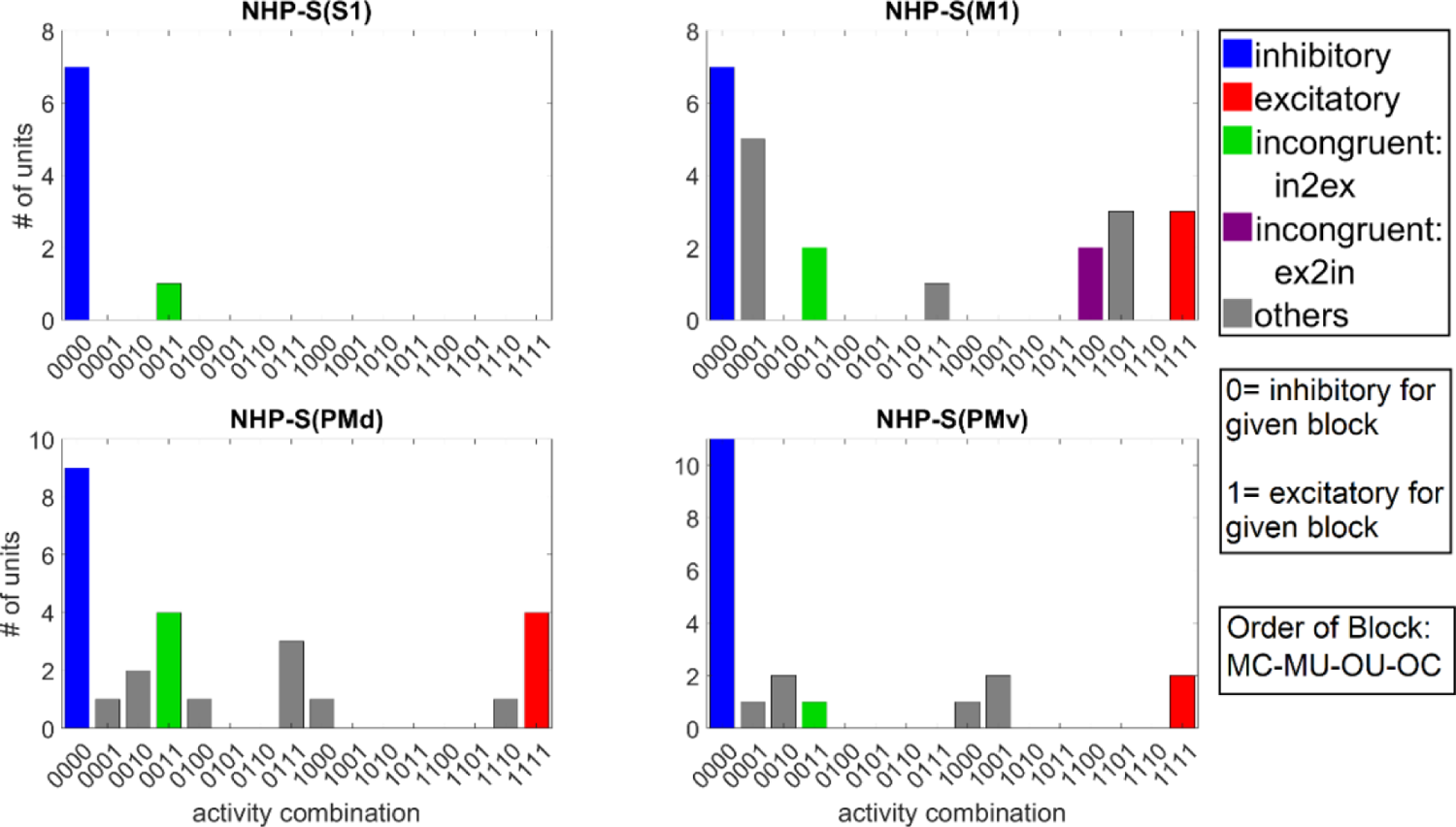
The Number of MN units from NHP S, for each possible activity combination during blocks in order of MC, MU, OU and OC. The y-axis represents the number of units and x-axis label shows the combination of inhibitory (0) or excitatory activity for the given four data blocks. The title on each subplot shows the corresponding cortex and NHP name.

**Figure S12:**
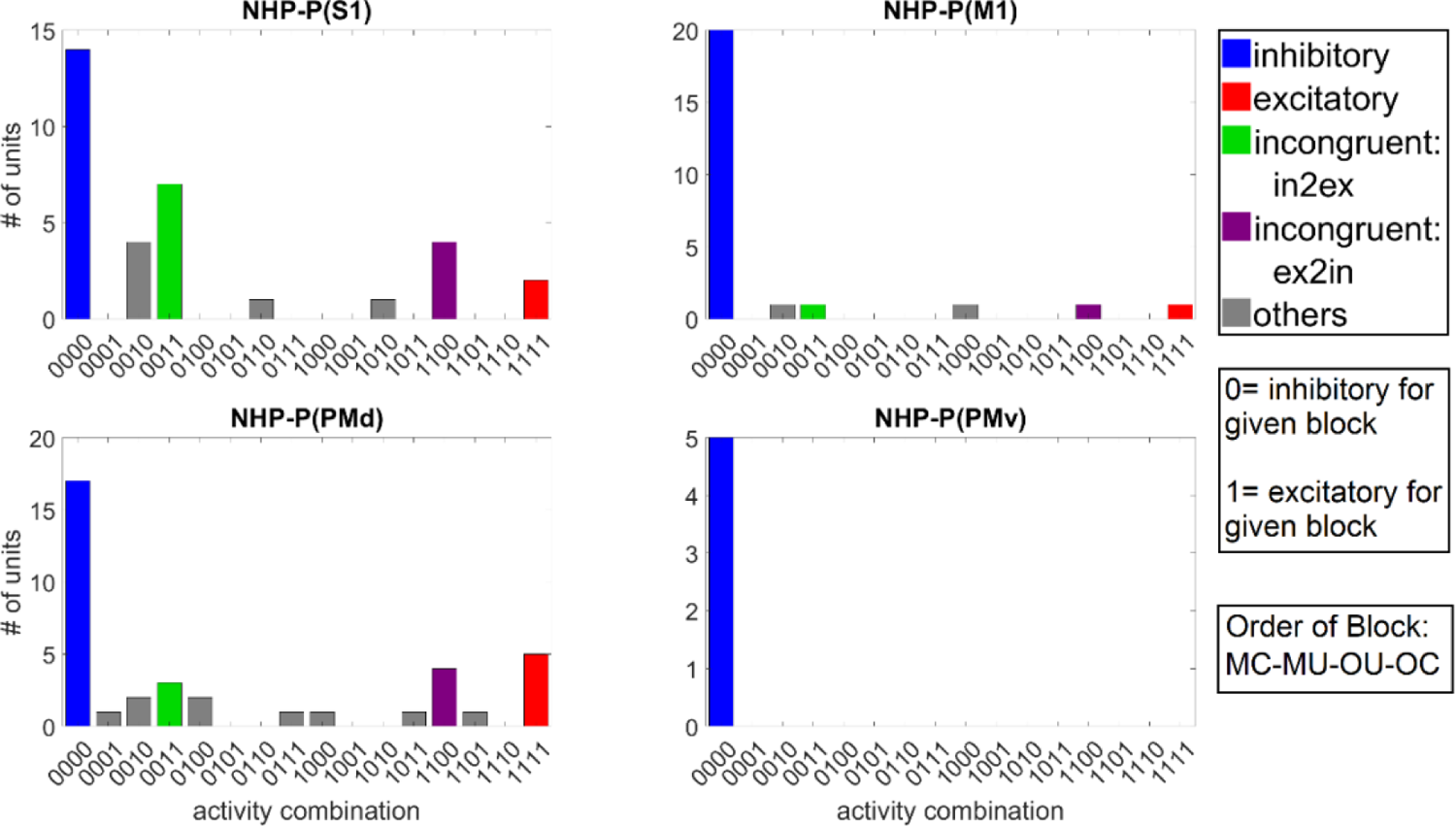
The Number of MN units from NHP P, for each possible activity combination during blocks in order of MC, MU, OU and OC. The y-axis represents the number of units and x-axis label shows the combination of inhibitory (0) or excitatory activity for the given four data blocks. The title on each subplot shows the corresponding cortex and NHP name.

**Figure S13:**
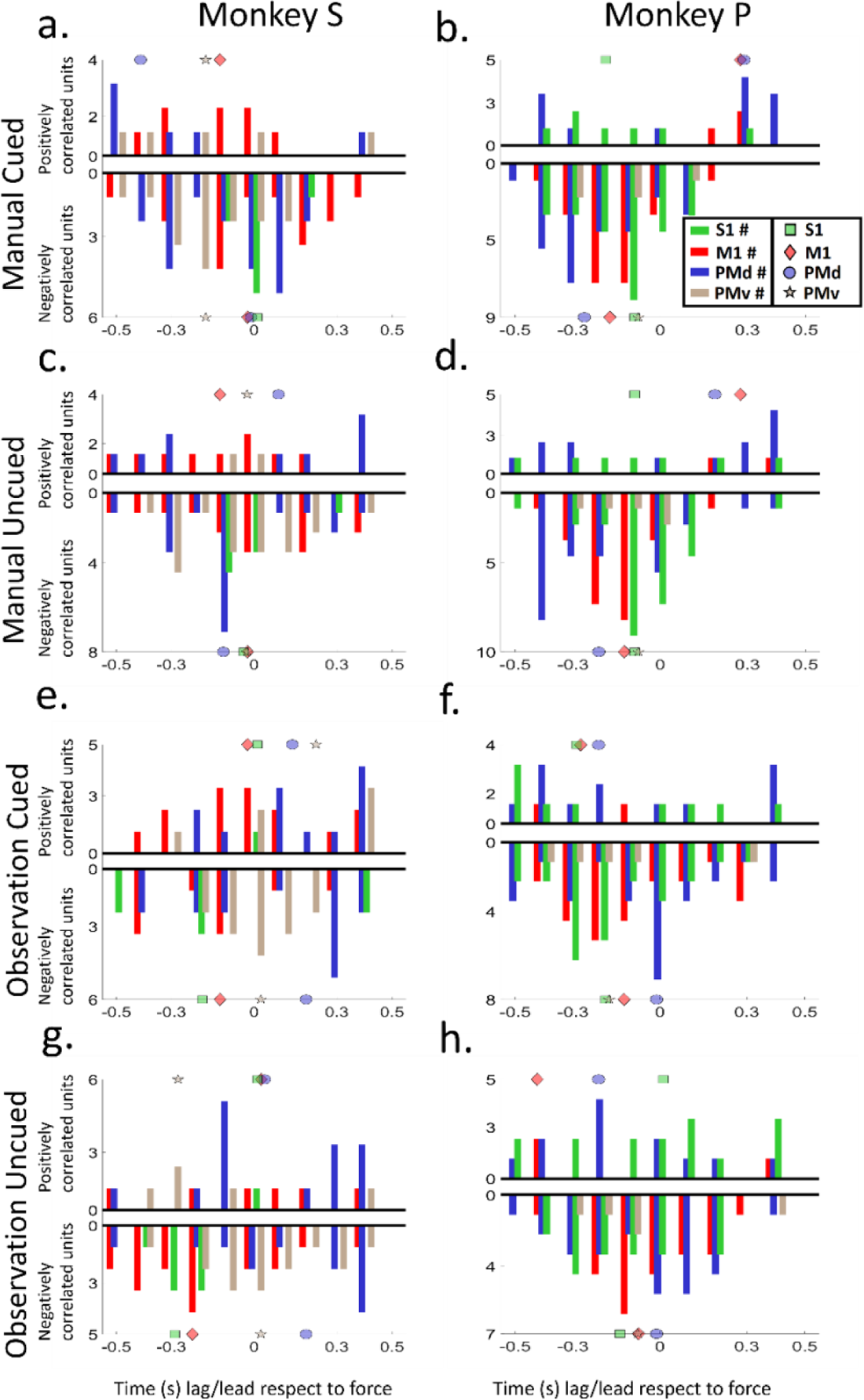
The number of units that had their highest correlation between force and spike rate for each time bin are shown leading (0.5s before) or lagging (0.5s after) each force value (100ms non-overlapping time bins). Units significant (F-test, p <0.05) on all block types were plotted in the time bin where their correlation coefficient with force was maximum in absolute value, either negative or positive. The y-axis represents the number of units (positively correlated units on the upper side and negatively correlated units on the lower). The x-axis is representing the time lag/lead for which the maximum correlation was found. Units from S1 (a, b), M1 (c, d), PMd (e, f) and PMv (g, h) cortex are given on each plot and the legend is showing the color diagram for each cortex.

**Figure S14:**
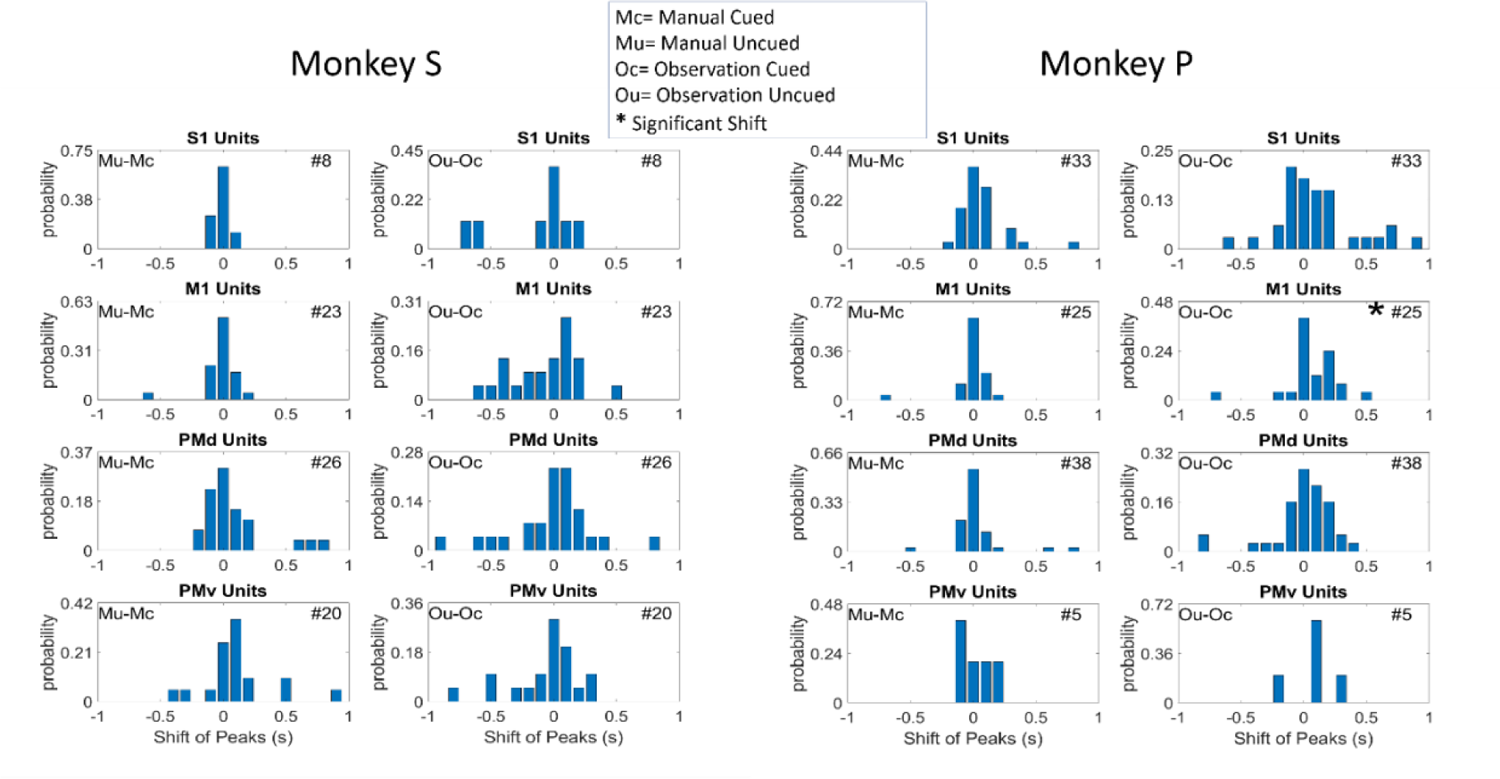
The shift of peak force correlation from manual to manual and observational to observational block. Bar plots showing the shift of peak correlation between Manual blocks and Observational blocks for S1, M1, PMd and PMv cortex for individual MNs. The shift from a block type to another block type is given on the left to of each figure and short form of the block type (Mc= Manual Cued, Mu= Manual Uncued, Oc= Observation Cued, Ou= Observation Uncued) was used. The number of units used is included on the upper right corner of each subplot. An asterisk (*) symbol before the number of units represents a significant shift in histogram between the blocks used for that subplot. Shift was calculated by subtracting the manual histogram for cued from the Manual histogram for uncued and similarly for the observational cases.

**Figure. S15:**
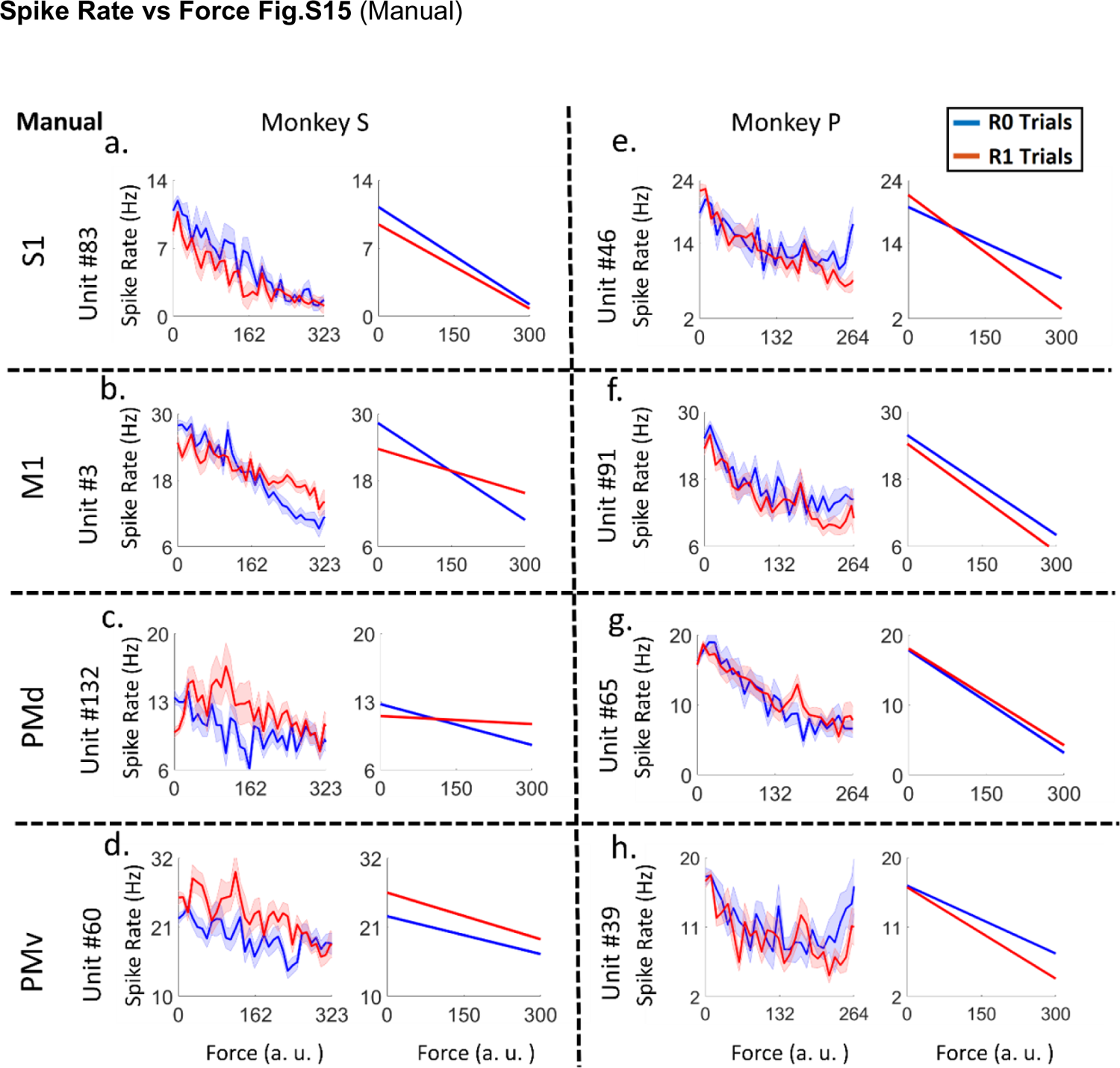
Plots of spike rate vs force (left subplots) and their linear tuning curve (right subplots) for Grip force manual trials as compared to Fig.9 for observational. The subplots show example units from S1 (plot *a, e*), M1 (plot *b, f*), PMd (plot *c, g*) and PMv (plot *d, h*) cortices of both NHPs (for NHP S plots *a, b, c* and *d* and for NHP P plots *e, f, g* and *h*). The units presented here showed a significant difference between R0 and R1 groups (ANCOVA, F-test, p<0.05) force tuning curves during observation of robotic manipulator applying grip force on a force sensor. Red lines indicate rewarding trials (R1) and blue indicate non-rewarding trials (R0).

**Figure S16:**
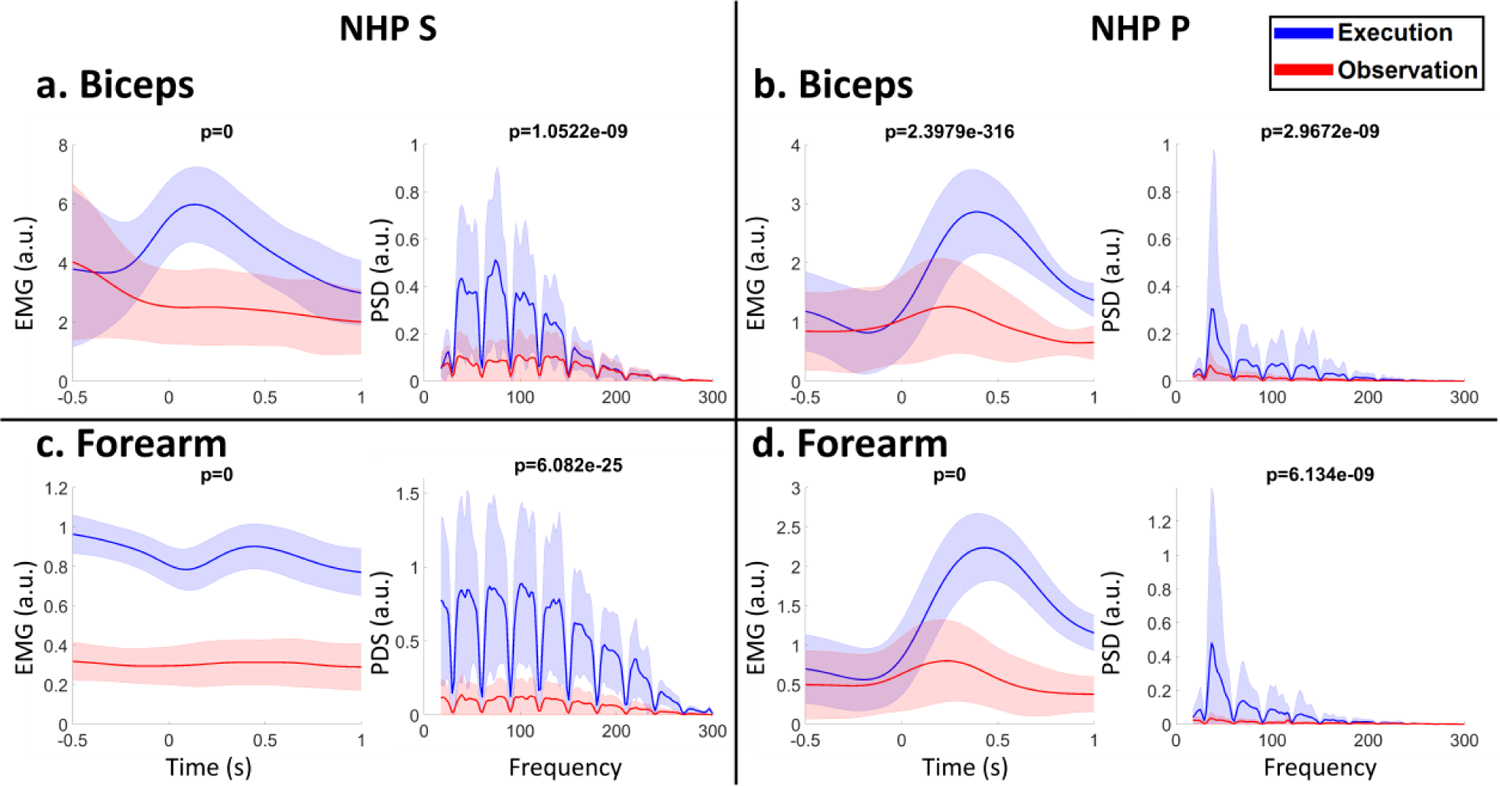
EMG collected from the Biceps (section a and b) and Forearm (section c and d) of NHP S (section a and c) and P (section b and d) recorded on different days. The blue color represents manual blocks (execution) and red is for observation. On each of four sections a, b, c and d, left plots show the mean EMG (0.5s pre-force onset to 1s post force onset activity) for all the trials with standard deviation as shaded error bar. The right plots are showing the mean power spectral density (PSD) with standard deviation for same data. A band pass filter was applied on the EMG for the power line noise and its harmonics (30Hz, 60Hz, 90Hz… etc) and the effect of that can be seen in the PSD plots (b). The p-value for the significance test between the mean manual and observational is given on the title of each subplot.

**Figure S17:**
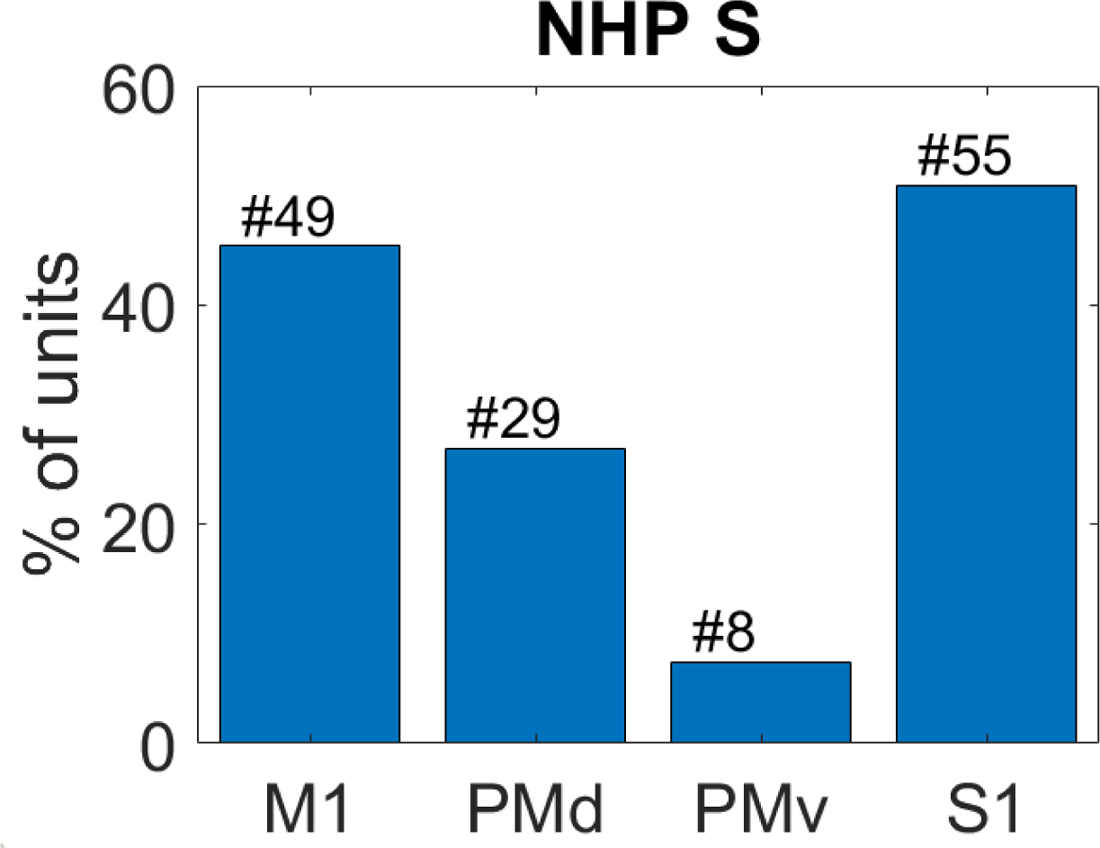
Percentage of with positive correlation between spike activity and extensor EMG envelope from the units that showed negative correlation with grip force.

**Figure S18:**
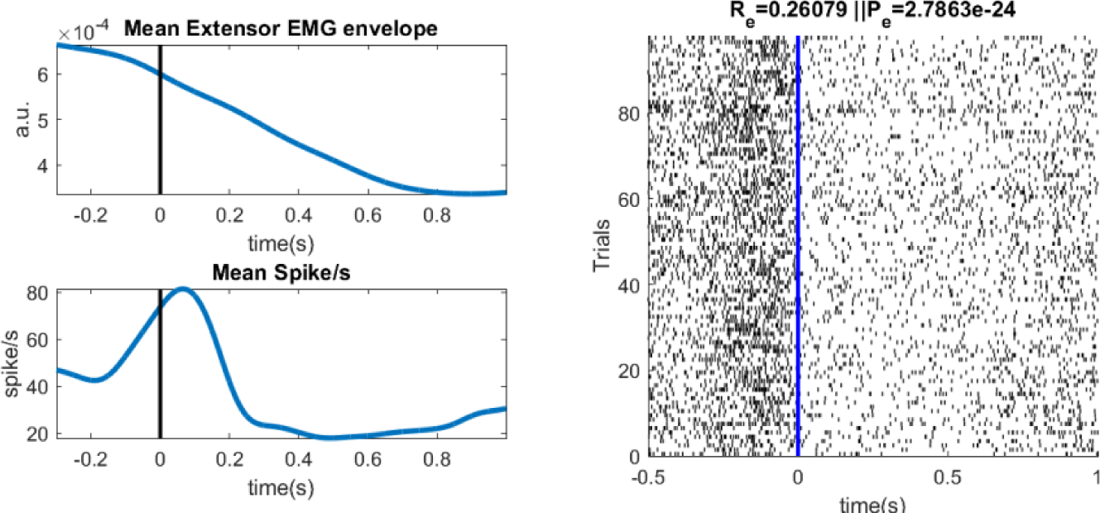
Example unit taken from NHP S, M1 cortex that shows positive correlation with the Extensor EMG activity. The left plots show the mean EMG envelope (top) and the mean spike activity (bottom) for all trials. The right raster plot shows spike information for all trials. The title on the raster pot is showing the spearman rank correlation (Re) with the significance (Pe) between binned (100ms) spike rate and extensor EMG done on concatenated data from all trials.

**Figure S19:**
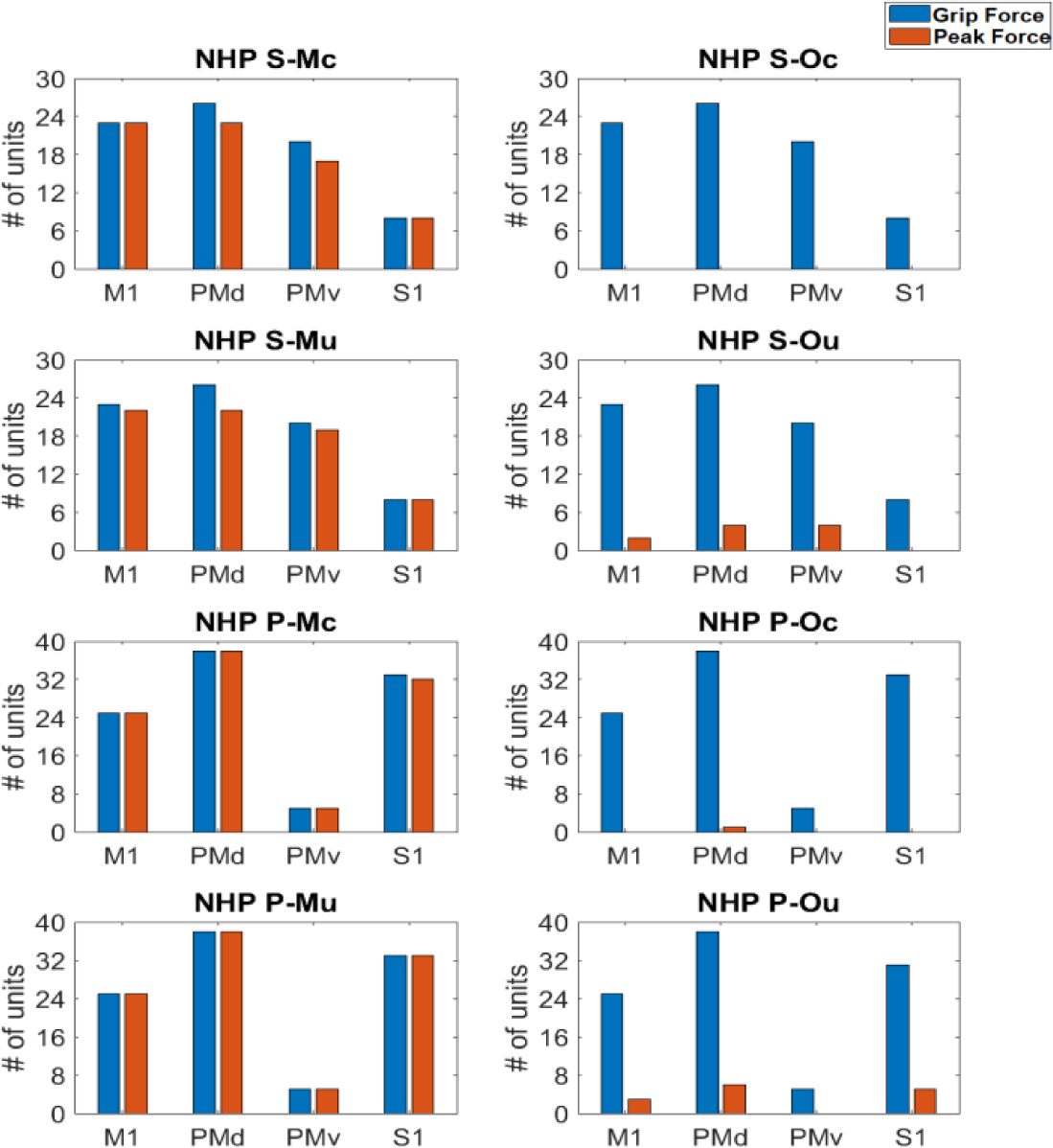
Significant mirror neurons (MN) for grip force (blue) and peak grip force (red) for Cued and uncued Manual (left subplots) and Observational (right subplots) block from NHP P (bottom four subplots) and S (top four subplots). The blue bars are representing units significant for grip force (F-test, BH corrected for number of population) and red bars are representing units significant for peak grip force values (F-test, BH corrected for number of population). The abbreviations on the subplot titles have the following meanings where Mc = Manual cued block, Oc = Observation cued block, Mu = Manual uncued block, and Ou = Observation uncued block. This plot is showing the mirror neuron units information from the Figure 6 of the main manuscript.

**Figure S20:**
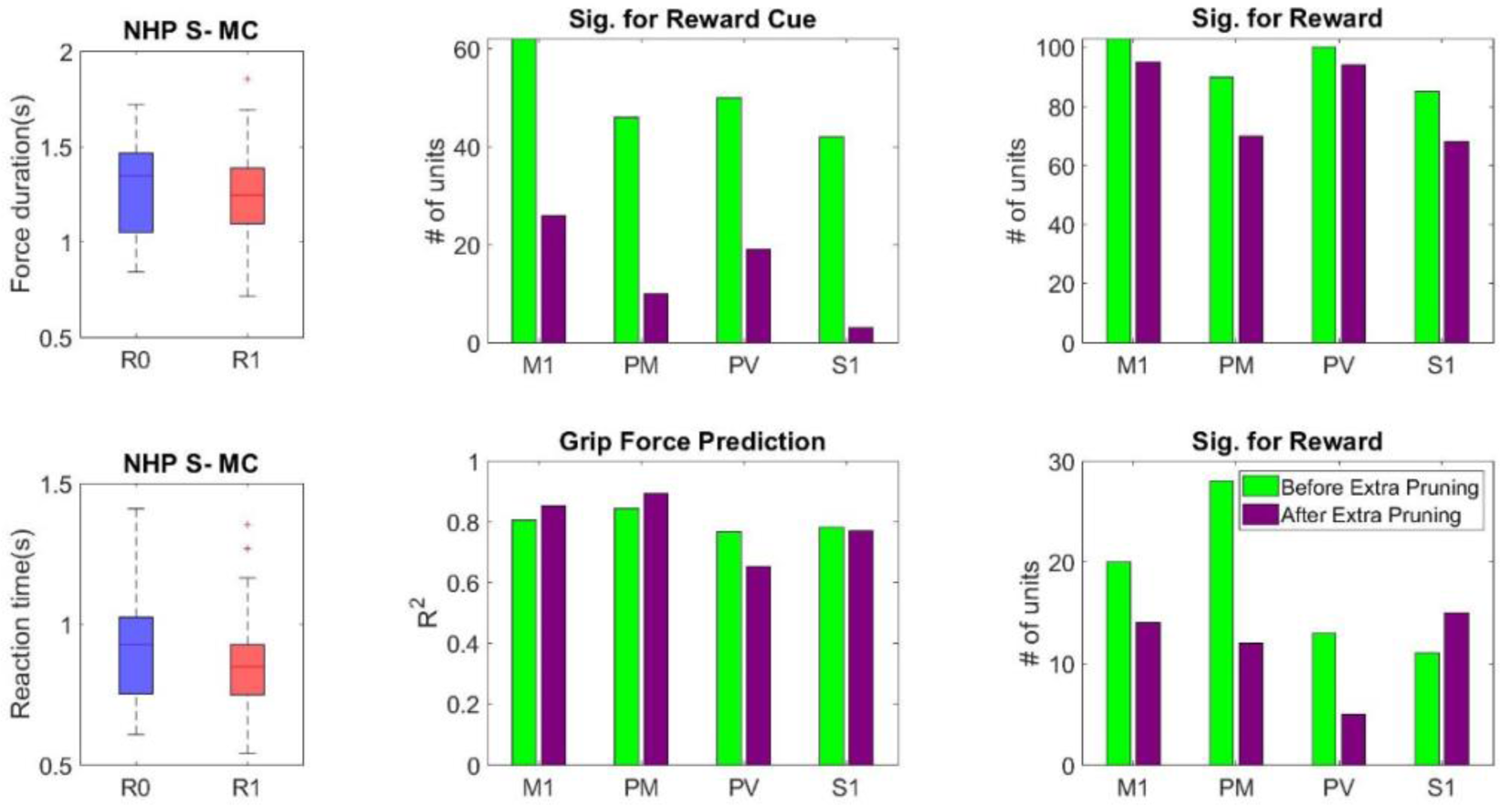
Statistical results from MC block of NHP S before and after pruning trials for the intention to use trials with similar motor behavior is given. The force duration and reaction time for R0 and R1 trials are given in left column respectively. The number of significant units for reward cue, reward grip force prediction and significant for reward outcome are indicated above the respective plots.

**Table S1:** Regression Model Output for NHP S & P for the two additional observational blocks recorded on a different day with units we could not track from the previous data sets.

| NHP S - S1 |  |  |  |  |  | NHP P - S1 |  |  |  |  |  |
| --- | --- | --- | --- | --- | --- | --- | --- | --- | --- | --- | --- |
| Block | R-fit | p-value | F-stat | R-predict | p-value | Block | R-fit | p-value | F-stat | R-predict | p-value |
| Observation Cued | 0.28 | 3e-91 | 3.81 | 0.19 | 8.3e-21 | Observation Cued | 0.46 | 1.3e-229 | 8.34 | 0.27 | 1.9e-26 |
| Observation Uncued | 0.34 | 4.3e-154 | 4.79 | 0.04 | 9.3e-6 | Observation Uncued | 0.53 | 0 | 10.09 | 0.13 | 1.8e-14 |
| NHP S - M1 |  |  |  |  |  | NHP P - M1 |  |  |  |  |  |
| Block | R-fit | p-value | F-stat | R-predict | p-value | Block | R-fit | p-value | F-stat | R-predict | p-value |
| Observation Cued | 0.65 | 0 | 11.59 | 0.6 | 2.6e-87 | Observation Cued | 0.61 | 0 | 13.19 | 0.55 | 2.7e-64 |
| Observation Uncued | 0.76 | 0 | 21.19 | 0.66 | 4.3e-113 | Observation Uncued | 0.69 | 0 | 18.07 | 0.55 | 3.1e-75 |
| NHP S - PMd |  |  |  |  |  | NHP P - PMd |  |  |  |  |  |
| Block | R-fit | p-value | F-stat | R-predict | p-value | Block | R-fit | p-value | F-stat | R-predict | p-value |
| Observation Cued | 0.53 | 1.9e-273 | 6.63 | 0.36 | 2.7e-43 | Observation Cued | 0.5 | 6.4e-274 | 10 | 0.52 | 6.3e-59 |
| Observation<br>Uncued | 0.67 | 0 | 11.57 | 0.39 | 0.55 | Observation<br>Uncued | 0.55 | 0 | 13.14 | 0.37 | 2e-44 |
| NHP S – PMv |  |  |  |  |  | NHP P – PMv |  |  |  |  |  |
| Block | R-fit | p-value | F-stat | R-predict | p-value | Block | R-fit | p-value | F-stat | R-predict | p-value |
| Observation<br>Cued | 0.6 | 0 | 11.84 | 0.72 | 5.7e-121 | Observation<br>Cued | 0.35 | 3.5e-197 | 11.24 | 0.28 | 4.9e-27 |
| Observation<br>Uncued | 0.8 | 0 | 34.11 | 0.72 | 6.1e-134 | Observation<br>Uncued | 0.46 | 0 | 13.18 | 0.36 | 1.4e-42 |

**Table S2:**
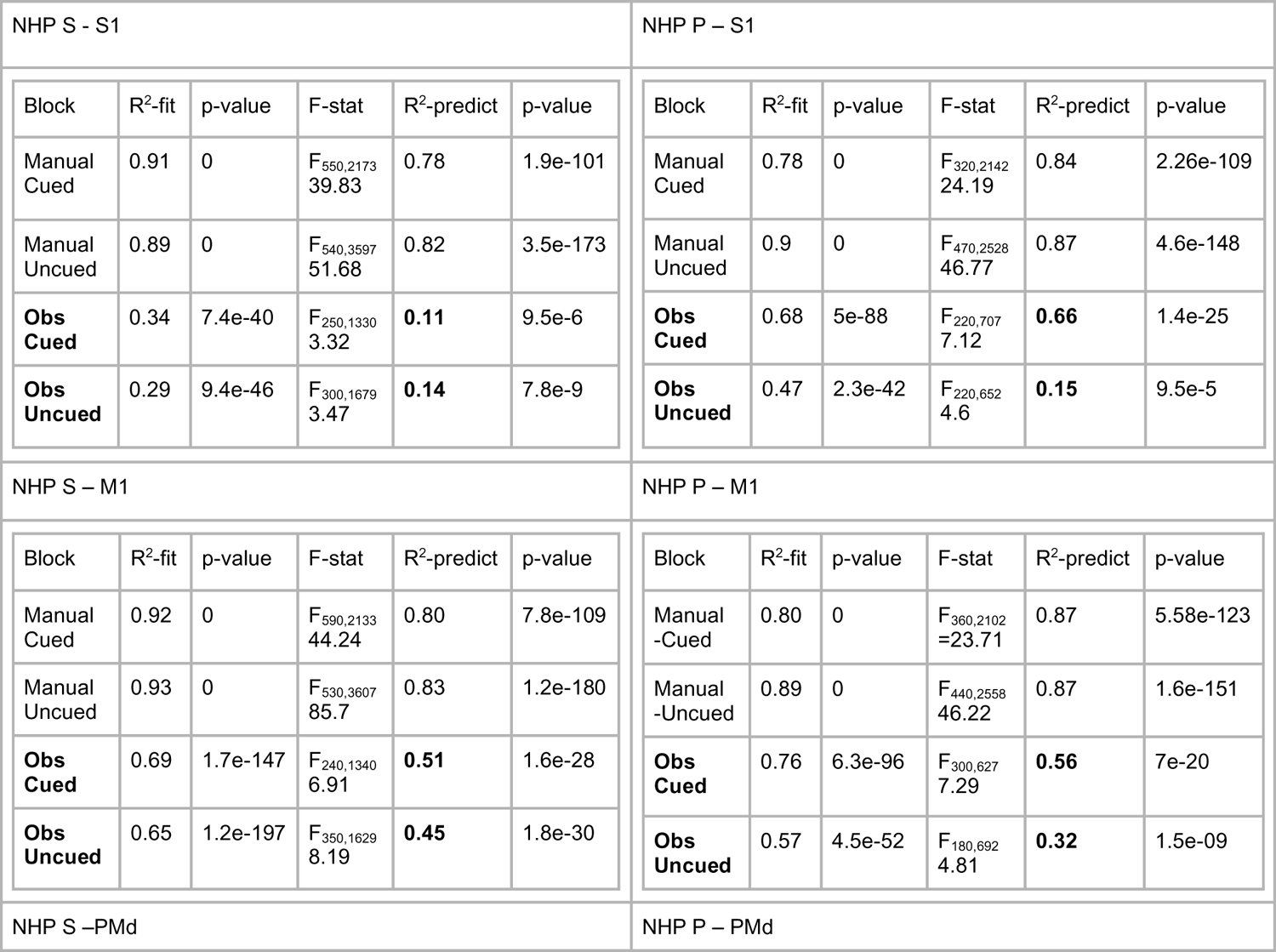
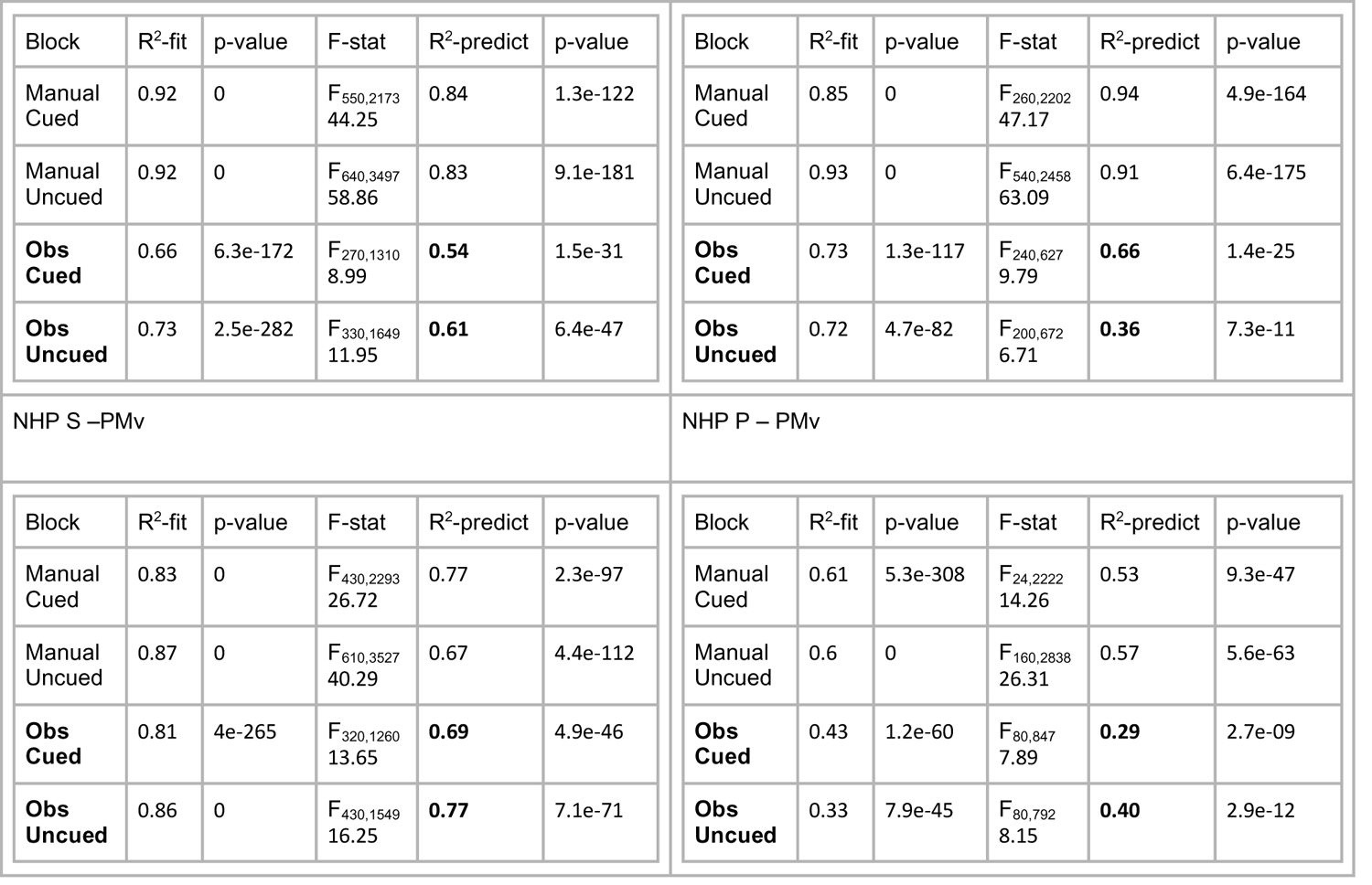
Results from the regression analysis for NHPs S and P between neural activity and force trajectories. The table contains the R-square value for the model (eq. 2) fit and prediction including their corresponding p-values and F-statistics. We have used bold text for the prediction during observation tasks. Note all single units were followed through each of the 4 task types on a given day. From table 1 it appears that these brain regions are representing force during both the manual trails and the observational versions of these trials. In Fig.6 we show the number of single units that significantly predict force during both manual and observational trails indicating that these same units are representing this information to some extent during both manual and observed movements, which may represent a new form of mirror neuron activity.

**Table S3:** Units with significantly different slopes for the force tuning curves between R0 and R1. The values in parenthesis show the adjusted significant units after Benjamini and Hochberg’s method accounting for the false discovery rate was applied. The BH method applied for number of units on which the hypothesis testing was applied in the population is described in the method section titled “grip force tuning curve analysis”.

|  | S1 # | S1 % | M1 # | M1 % | PMd # | PMd % | PMv # | PMv % |
| --- | --- | --- | --- | --- | --- | --- | --- | --- |
| NHP S Manual | 11 (3) | 8 (2) | 20 (4) | 14 (3) | 28 (24) | 20 (17) | 13 (4) | 8 (2) |
| NHP P Manual | 9 (1) | 7 (1) | 9 (4) | 8 (3) | 16 (6) | 14 (5) | 4 (0) | 5 (0) |
| NHP S Obs. | 2 (2) | 2 (2) | 2 (1) | 2 (1) | 2 (0) | 2 (0) | 0 (0) | 0 (0) |
| NHP P Obs. | 0 (0) | 0 (0) | 1 (0) | 1 (0) | 2 (0) | 2 (0) | 1 (1) | 2 (2) |

**Table S4:** Units with significantly different y-intercept for the force tuning curves for R0 and R1. The values in parenthesis show the adjusted significant units after Benjamini and Hochberg’s method accounting for the false discovery rate was applied. The BH method applied for number of population is described in the method section titled “grip force tuning curve analysis”.

|  | S1 # | S1 % | M1 # | M1 % | PMd # | PMd % | PMv # | PMv % |
| --- | --- | --- | --- | --- | --- | --- | --- | --- |
| NHP S Manual | 14 (2) | 10 (1) | 29 (21) | 21 (15) | 32 (24) | 23 (17) | 27 (17) | 16 (10) |
| NHP P Manual | 8 (0) | 6 (0) | 7 (1) | 6 (1) | 14 (4) | 12 (3) | 2 (0) | 2 (0) |
| NHP S Obs. | 3 (3) | 3 (3) | 4 (4) | 3 (3) | 1 (1) | 1 (1) | 4 (4) | 3 (3) |
| NHP P Obs. | 0 (0) | 1 (0) | 1 (0) | 1 (0) | 1 (0) | 1 (0) | 2 (0) | 3 (0) |

**Table S5:** the p-value from the Rayleigh test.

|  | S1 | M1 | PMd | PMv |
| --- | --- | --- | --- | --- |
| NHP S | 0.234 | 0.0036 | 0.7366 | 0.0595 |
| NHP P | 0.4617 | 0.0045 | 0.4104 |  |
| Combined | 0.5714 | 1.48E-05 | 0.3726 |  |

## Notes

**Acknowledgements**: We thank Matthew J. Perry for helping with the manuscript. Research was supported by NIH 1R01NS092894-01, NSF IIS-17 1527558, DARPA REPAIR Project N66001-10-C-2008.

### Competing Interest Statement

The authors have declared no competing interest.

### Summary of Updates

The update is too extensive to list.

